# Heme drives cardiac endothelial senescence in sepsis via STING activation

**DOI:** 10.1101/2025.06.18.660159

**Authors:** Tingting Li, Peilin Zhu, Tao Zhang, Joseph Adams, Fei Tu, Chloe Garbe, Xiaojin Zhang, Li Liu, Krishna Singh, David L Willams, Chuanfu Li, Xiaohui Wang

**Author notes:** These authors contributed equally to this study (Tingting Li, Peilin Zhu). Corresponding author: Xiaohui Wang, Ph.D. Assistant Professor Department of Biomedical Sciences East Tennessee State University Johnson City, TN 37614.

## Abstract

Sepsis-induced cardiac dysfunction is a major contributor to sepsis-related mortality, and many patients continue to experience long-term cardiac complications after recovery. Here, we demonstrate that cardiac senescence is a key feature of sepsis-associated cardiac dysfunction, with endothelial cells identified as the predominant senescent population in septic cardiac tissue. Yet, the pathogenic drivers of endothelial senescence in sepsis remain poorly characterized. Among potential mediators, we found that elevated levels of heme, a byproduct of hemolysis, strongly correlate with increased endothelial senescence and impaired cardiac function. Mechanistic studies revealed that heme acts as a novel ligand for STING, exacerbating bacterial infection induced STING polymerization and activation, thereby promoting endothelial senescence. Notably, either STING inhibition or enhanced heme clearance via increased hemopexin expression significantly alleviated cardiac endothelial senescence and facilitated cardiac functional recovery in septic mice. These findings identify heme as a critical pathogenic driver of endothelial senescence and highlight heme clearance as a promising therapeutic strategy for mitigating sepsis induced cardiac dysfunction.

## Introduction

Sepsis, a life-threatening condition characterized by a dysregulated immune response to infection, remains a leading cause of mortality in intensive care units worldwide, accounting for approximately 20% of all global deaths^1, 2^. Among the various organs affected in sepsis, the heart is especially susceptible to injury^3, 4, 5^. Sepsis-induced cardiomyopathy is a major contributor to poor outcomes, with many septic patients experience long-term cardiac complications. However, its underlying pathophysiological mechanisms remain poorly defined, and no targeted therapies are currently available.

Endothelial cells, which constitute the largest organ in the body, are essential for maintaining vascular integrity, regulating immune responses, and supporting metabolic homeostasis^6, 7, 8^. Endothelial dysfunction is a hallmark of sepsis and a critical driver of disease progression^9, 10, 11,12^. In the heart, endothelial injury disrupts tissue oxygenation, increases vascular leakage, and exacerbates inflammation, thereby contributing to myocardial dysfunction^10, 12^. Although mechanisms such as cytokine release, oxidative stress, coagulopathy, and glycocalyx degradation have been implicated in septic endothelial injury^11, 12, 13^, the factors driving cardiac endothelial dysfunction during sepsis remain largely undefined.

Another important yet underexplored mechanism that may contribute to septic endothelial injury is cellular senescence. Cellular senescence, characterized as a state of cell cycle arrest and the acquisition of a proinflammatory senescence-associated secretory phenotype (SASP), has emerged as a critical contributor to endothelial dysfunction in various cardiovascular diseases^14, 15, 16, 17, 18^. Senescent endothelial cells lose their regenerative capacity, promote chronic inflammation, and impair vascular repair^15, 16, 19, 20, 21^. Although recent studies suggest that sepsis is associated with endothelial cell loss and reduced proliferative capacity, it remains unclear whether sepsis actively induces endothelial senescence and whether this process contributes to the development of sepsis-associated cardiomyopathy and impairs cardiac functional recovery following sepsis.

Hemolysis is a common feature of sepsis and is strongly associated with disease severity and poor clinical outcomes^22, 23, 24^. During hemolysis, the breakdown of red blood cells leads to the release of free heme into the circulation^25^. Free heme is now recognized as a potent danger-associated molecular pattern (DAMP) that promotes oxidative stress, amplifies inflammation, and induces cell death^22, 24, 26, 27, 28, 29^. However, whether free heme contributes to endothelial dysfunction and cardiac impairment during sepsis remains unclear.

In the present study, we identify pronounced cardiac endothelial senescence as a hallmark of sepsis. Senescence levels correlated with disease severity and were associated with impaired vascular repair capacity and adverse outcomes. We further demonstrate that free heme, released during hemolysis in sepsis, acts as a key mediator of cardiac endothelial senescence and dysfunction. Elevated heme levels are strongly associated with impaired cardiac function and increased mortality. Although our recent work showed that heme contributes to Kupffer cell senescence via cGAS-STING activation, the mechanisms by which heme activates this pathway remained unclear^22^. In this study, we provide mechanistic evidence that heme functions as a novel ligand of STING, promoting its oligomerization and subsequently amplifying STING-TBK1 activation, a known trigger of endothelial senescence^30, 31, 32^. Importantly, pharmacological inhibition of STING effectively mitigated endothelial senescence and improved cardiac functional recovery in septic mice. Furthermore, therapeutic upregulation of hemopexin, a heme scavenger^22, 23^, significantly reduced circulating free heme levels, alleviated cardiac endothelial senescence, and enhanced cardiac functional recovery during sepsis.

Collectively, these findings uncover a novel role for heme in driving cGAS–STING mediated endothelial senescence and cardiac dysfunction during sepsis and highlight heme clearance as a promising therapeutic strategy to improve sepsis outcomes.

## Results

### 1. Sepsis induces cardiac senescence that correlates with disease severity

Given the critical role of cellular senescence in cardiovascular pathology and its potential contribution to sepsis induced cardiac dysfunction^16, 33, 34, 35^, we first examined whether sepsis promotes cardiac senescence. Sepsis was induced in mice using the cecal ligation and puncture (CLP) model, and heart tissues were collected six hours post-sepsis for analysis.

Gene set enrichment analysis (GSEA) of RNA sequencing (RNA-seq) data revealed a significant upregulation of gene signatures associated with cellular senescence in septic hearts (*Figures 1A-C*). Notably, senescence associated secretory phenotype (SASP) markers, including MMPs, IL-1α, IL-1β, and Cxcl family cytokines, as well as the cell cycle inhibitors p21 and p16, were markedly increased in septic hearts compared to sham controls (*Figures 1A-C&S1C*). Consistent with transcriptomic findings, western blot analysis showed a substantial increase in protein levels of key senescence markers, including p21, p16, and acetylated p53 (Ac-p53), in heart tissues from septic mice (*Figure 1D*). To investigate whether cardiac senescence correlates with sepsis severity, we compared the expression of senescence markers between survivors and non-survivors. Since a body temperature below 30°C is a reliable predictor of mortality in septic mice, we stratified mice at 24 hours post-CLP into survivors (body temperature >30°C) and non-survivors (body temperature <30°C). qPCR analysis revealed significantly higher expression of p21, p16, and SASP factors in non-survivors compared with survivors (*Figures 1E-H&S1B*). Together, these results provide compelling evidence that sepsis induces severe cardiac senescence, which correlates with septic severity.

**Figure. 1.**
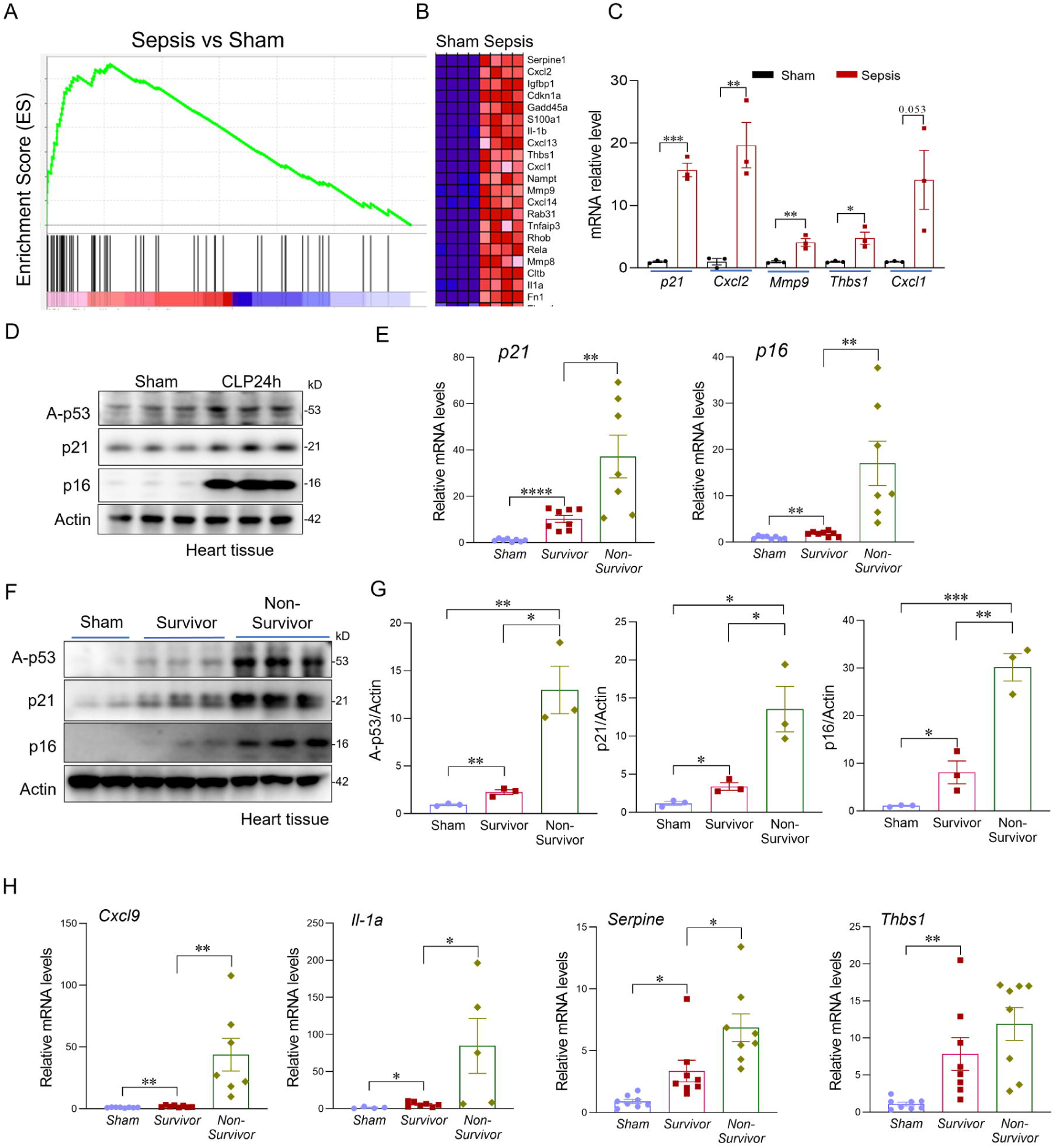
Sepsis induces cardiac senescence that correlates with disease severity. (**A**) Gene set enrichment analysis (GSEA) of RNA-seq data from hearts of sham and septic mice, highlighting enrichment of senescence-related genes. (**B**) Heat map showing differential expression of selected senescence-associated genes. (**C**) Expression levels of representative senescence-related genes (Cdkn1a/p21, Cxcl2, Mmp9, Thbs1, Cxcl1) based on RNA-seq analysis of hearts from sham and septic mice. (**D**) Western blot analysis of senescence markers p16, p21, and acetylated p53 in sham and septic heart tissues (n=6 mice/group). (**E**) qRT-PCR analysis of senescence markers in sham and septic heart tissues (n=6-8 mice/group). (**F, G**) Western blot analysis and quantification of senescence markers p16, p21, and acetylated p53 in heart tissues from sham mice, septic non-survivors (body temperature < 30L°C), and septic survivors (> 30L°C) at 24 hours post-CLP (n=6 mice/group). (**H)** qRT-PCR analysis of senescence associated genes (Cxcl9, Il-1A, Serpine1, Thbs1) in heart tissues from sham mice, septic survivors, and septic non-survivors. All data are presented as mean ± SD. Statistical significance was determined using the unpaired two-tailed Student’s t-test: *PL<L0.05, **PL<L0.01, ***PL<L0.001, ****PL<L0.0001.

### 2. Cardiac endothelial cells are the predominant senescent population during sepsis

To identify the primary senescent cell population in the septic heart, we performed co-immunostaining of the senescence markers p21 and p16 with CD31, an established endothelial cell marker. Quantitative analysis showed a ∼ 30 fold increase in p21 positive endothelial cells and a ∼16 fold increase in p16 positive endothelial cells in septic heart tissues compared to sham controls (*Figures 2A-D*). Furthermore, p21 and p16 staining largely co-localized with CD31 (*Figures 2A-D*), indicating that endothelial cells are the predominant senescent population in the septic heart. To investigate whether cardiac endothelial senescence correlates with sepsis severity, we compared senescence marker expression between survivors and non-survivors at 24 hours post-CLP. As shown in Figures S1A-D, non-survivors exhibited significantly higher levels of endothelial senescence, as evidenced by an increased number of p21 and p16 positive endothelial cells, compared to survivors (*Figures S1A-D*). Because cellular senescence impairs endothelial cell proliferation, which is essential for vascular repair following injury, we next assessed endothelial proliferation in both survivors and non survivors. Immunostaining for Ki67, a marker of actively dividing cells, revealed that endothelial proliferation was markedly reduced in non-survivors (*Figures 2E-F&S1E-F*).

**Figure. 2.**
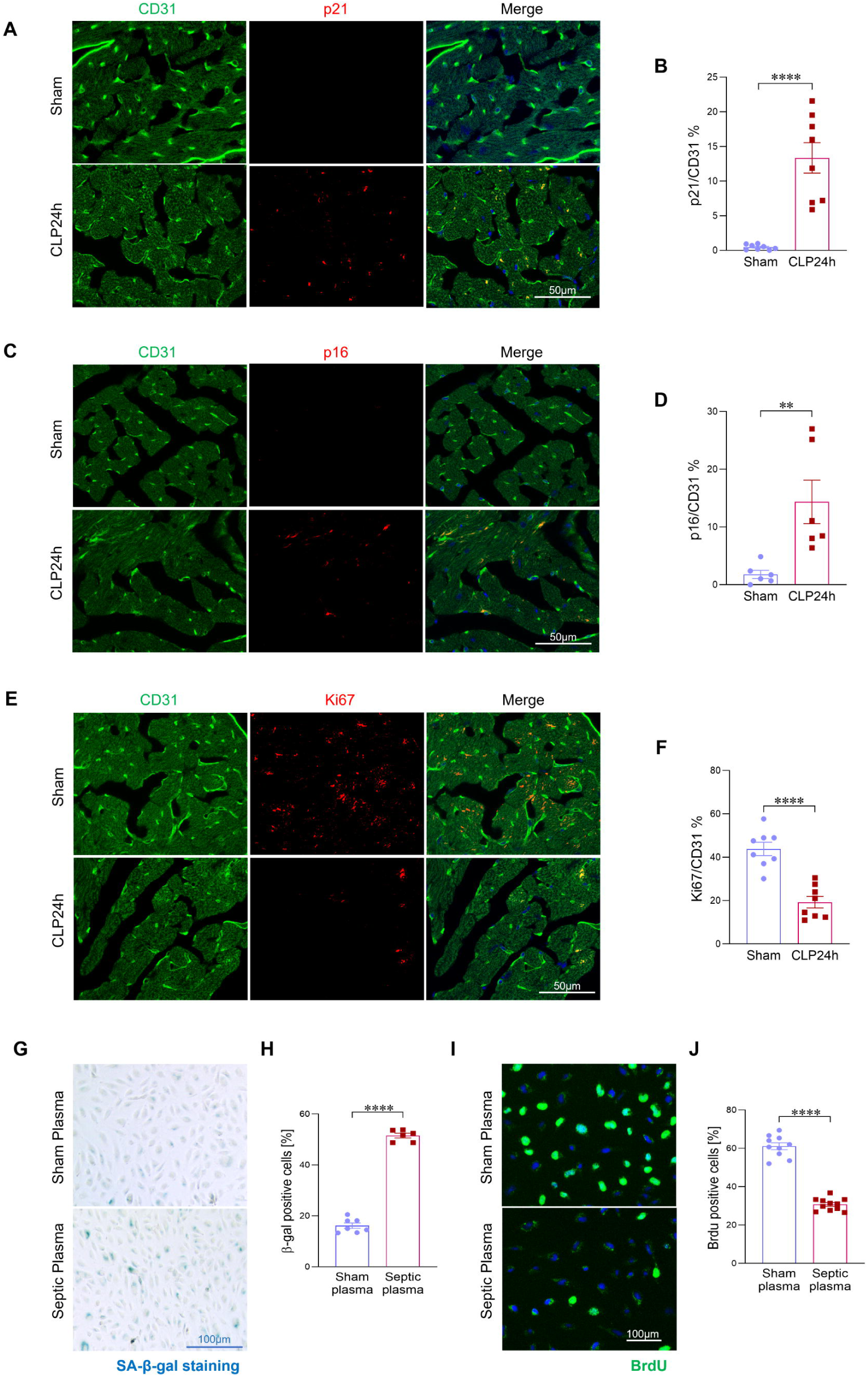
**Cardiac endothelial cells are the predominant senescent population during sepsis**. (**A-D**) Co-immunostaining of senescence markers p21 and p16 with the endothelial cell marker CD31, and their quantification in heart sections from sham and septic mice (n=6 mice/group). Scale bars, 50Lμm. (**E-F**) Co-immunostaining of the cellular proliferation marker Ki67 with the endothelial marker CD31 in heart tissues from sham controls and septic mice at 24 hours post-CLP, along with corresponding quantification. (n=8/group). Scale bars, 50Lμm. (**G**) β-gal staining of HUVECs treated with plasma (1:200) from sham or septic (CLP 24 h) mice for 24 hours (n=6/group). Scale bars, 100Lμm. (**H**) Quantification of β-gal staining in HUVECs, as shown in (**G**). (**I**) BrdU incorporation assay in HUVECs from the indicated treatment groups (n=6/group). Scale bars, 100Lμm. (**J)** Quantification of BrdU incorporation in HUVECs, as shown in (**H**). All data are presented as mean ± SD. Statistical analysis was performed using the unpaired two-tailed Student’s t-test: *PL<L0.05, **PL<L0.01, ****PL<L0.0001.

**Figure S1.**
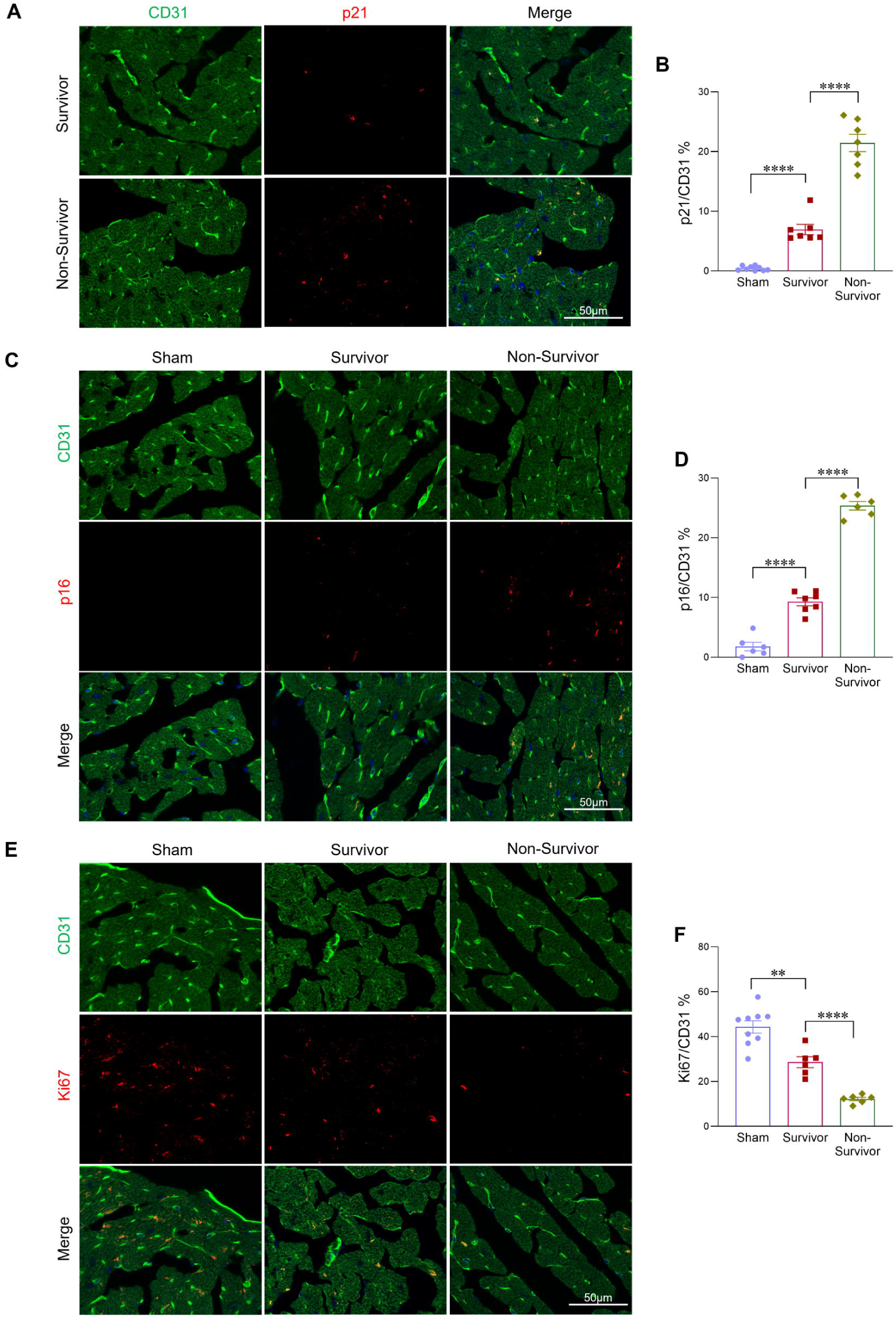
**Cardiac endothelial senescence impairs proliferation in sepsis**. (**A-B**) Immunofluorescent co-staining and quantification of the senescence marker p21 (red) and the endothelial marker CD31 (green) in heart sections from the indicated groups (n=6 mice/group). Scale bars, 50Lμm. (**C-D**) Immunofluorescent co-staining and quantification of the senescence marker p16 (red) and the endothelial marker CD31 (green) in heart sections from the indicated groups (n=6 mice/group). Scale bars, 50Lμm. (**E, F**) Immunofluorescent co-staining and quantification of the proliferation marker Ki67 (red) and the endothelial marker CD31 (green) in heart sections from sham, septic survivors, and septic non-survivors (n=6-8 mice/group). Scale bars, 50Lμm. All data are presented as mean ± SD. Statistical analysis was performed using the unpaired two-tailed Student’s t-test: *PL<L0.05, **PL<L0.01, ****PL<L0.0001.

To further determine whether circulating factors in sepsis can directly promote endothelial cell senescence and inhibit proliferation, we exposed human umbilical vein endothelial cells (HUVECs) to plasma collected from either septic or sham mice for a duration of 24 hours. Cells treated with septic plasma showed a significantly greater number of β-galactosidase positive cells compared to controls, indicating the presence of senescence-inducing factors in the plasma (*Figures 2G-H*). In parallel, BrdU incorporation assays revealed a significant reduction in cell proliferation following exposure to septic plasma, further supporting the notion that sepsis-associated circulating mediators directly compromise endothelial cell proliferation (*Figures 2I-J*). Together, these findings demonstrate that cardiac endothelial cells are the main senescent population in sepsis, showing high susceptibility to systemic inflammatory stress and impaired proliferation. This dysfunction may compromise vascular repair and contribute to organ damage, highlighting endothelial senescence as a potential therapeutic target in sepsis.

### 3. Elevated heme levels correlate with cardiac endothelial senescence and dysfunction in sepsis

Hemolysis and the subsequent release of free heme are well recognized complications in sepsis^22, 23, 25^. Considering the association between elevated heme levels and sepsis severity, we investigated whether circulating heme contributes to sepsis induced endothelial cell senescence. To assess this, we first measured circulating free heme levels and examined their correlation with cardiac endothelial senescence in septic mice (*Figures 3A-C*). As shown in Figure. 3B-C, septic mice with higher plasma heme concentrations exhibited a marked increase in endothelial cell senescence compared to those with lower heme levels. This finding reveals a positive correlation between circulating heme levels and the severity of cardiac endothelial senescence. To further elucidate the causal role of heme in endothelial senescence during sepsis, we administered exogenous heme (15 mg/kg, IV injection) or vehicle control (PBS) immediately following CLP-induced sepsis. At 24 hours post-CLP, Heme treated mice displayed significantly elevated endothelial senescence in the heart compared to vehicle treated controls (*Figures 3D-E, S2A-B*). In addition, echocardiographic analysis demonstrated that exogenous heme administration let to a significant reduction in cardiac function, as evidenced by decreased ejection fraction (EF) and fractional shortening (FS) (*Figures 3F-H*). Western blot analysis further revealed significantly increased expression of senescence markers, including acetyl-p53, p21, and p16, in septic heart tissues treated with exogenous heme compared to control septic tissues (*Figures 3I-J*). Collectively, these results suggest that elevated circulating heme not only correlates with but also exacerbates cardiac endothelial senescence and contributes to impaired cardiac function during sepsis.

**Figure. 3.**
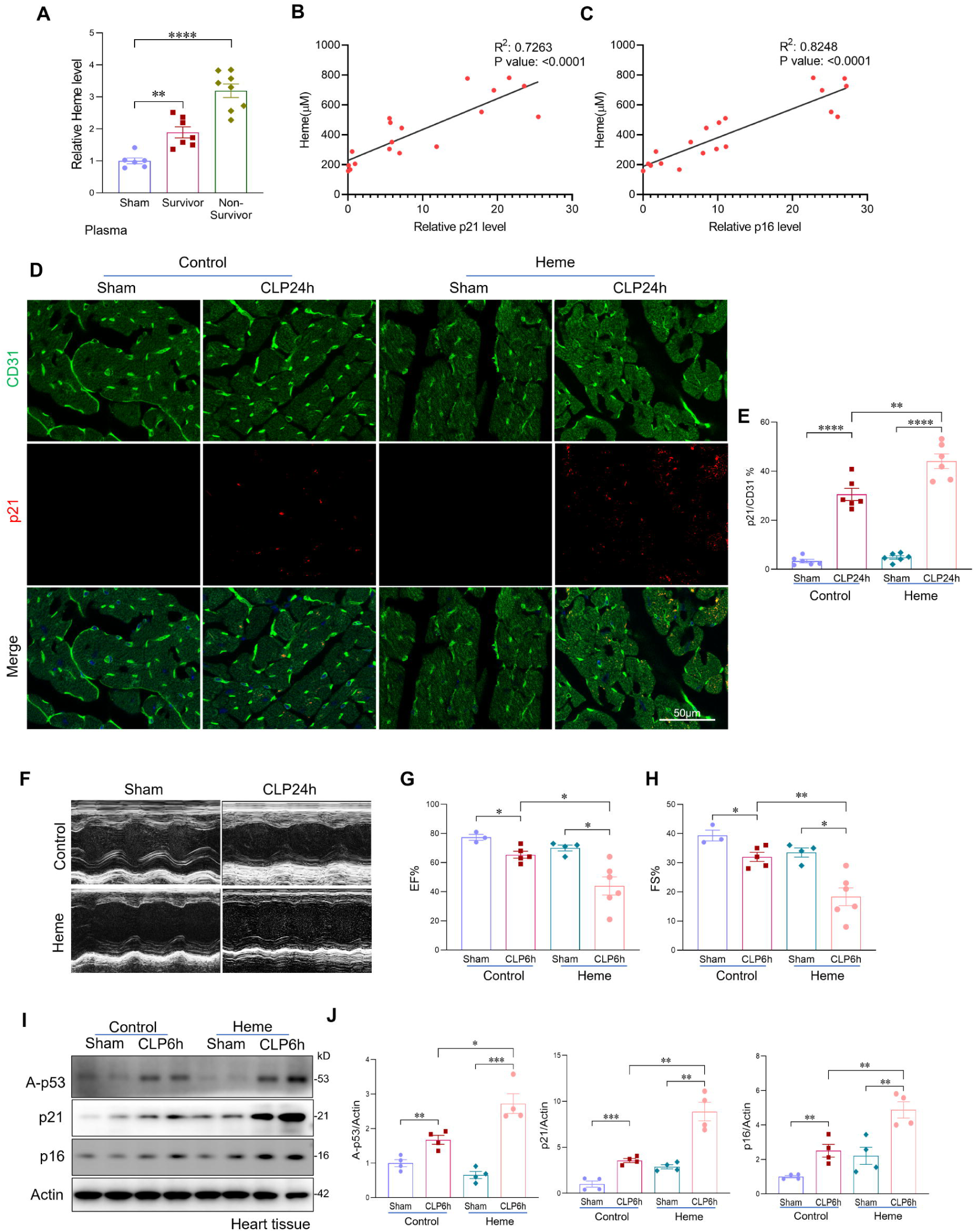
**Elevated heme levels correlate with cardiac endothelial senescence and dysfunction in sepsis**. (**A**) Plasma heme levels in sham mice, septic non-survivors (body temperature < 30L°C), and septic survivors (> 30L°C) at 24 hours post-CLP (n=6–8 mice/group). (**B, C**) Mice with higher circulating heme levels exhibited increased cardiac endothelial senescence, as indicated by elevated expression of senescence markers p21 and p16. (**D, E**) Co-immunostaining of p21 with endothelial cells marker CD31 and quantification in heart tissues from sham and septic mice with or without heme treatment (n=6 mice/group). Scale bars, 50Lμm. (**F–H**) Echocardiographic assessment of ejection fraction (EF) and fractional shortening (FS) in sham and septic mice treated with or without heme (n=6 mice/group). (**I, J**) Western blot analysis and quantification of senescence markers p16, p21, and acetylated p53 in heart tissues from sham and septic mice with or without heme treatment (n=6 mice/group). All data are presented as mean ± SD. Statistical analysis was performed using the unpaired two-tailed Student’s t-test: *PL<L0.05, **PL<L0.01, ***PL<L0.001, ****PL<L0.0001.

**Figure S2.**
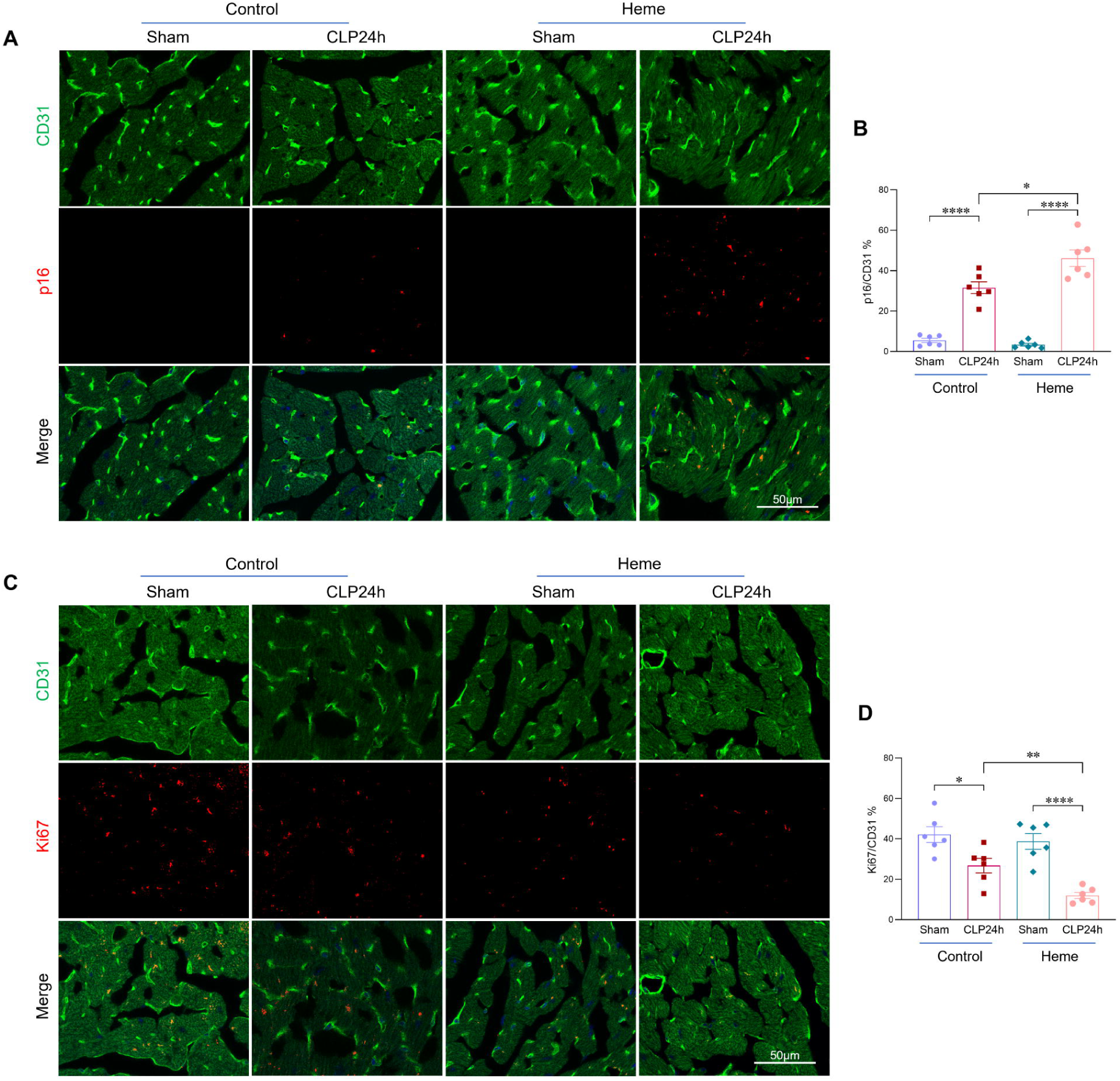
**Heme administration enhances cardiac endothelial senescence and impairs endothelial proliferation**. (**A, B**) Immunofluorescent co-staining and quantification of the senescence marker p16 (red) and the endothelial marker CD31 (green) in heart sections from the indicated groups (n=6 mice/group). Scale bars, 50Lμm. (**C, D**) Immunofluorescent co-staining and quantification of the proliferation marker Ki67 (red) and the endothelial marker CD31 (green) in heart sections from the indicated groups (n=6 mice/group). Scale bars, 50Lμm. All data are presented as mean ± SD. Statistical analysis was performed using the unpaired two-tailed Student’s t-test: *PL<L0.05, **PL<L0.01, ****PL<L0.0001.

**Figure S3.**
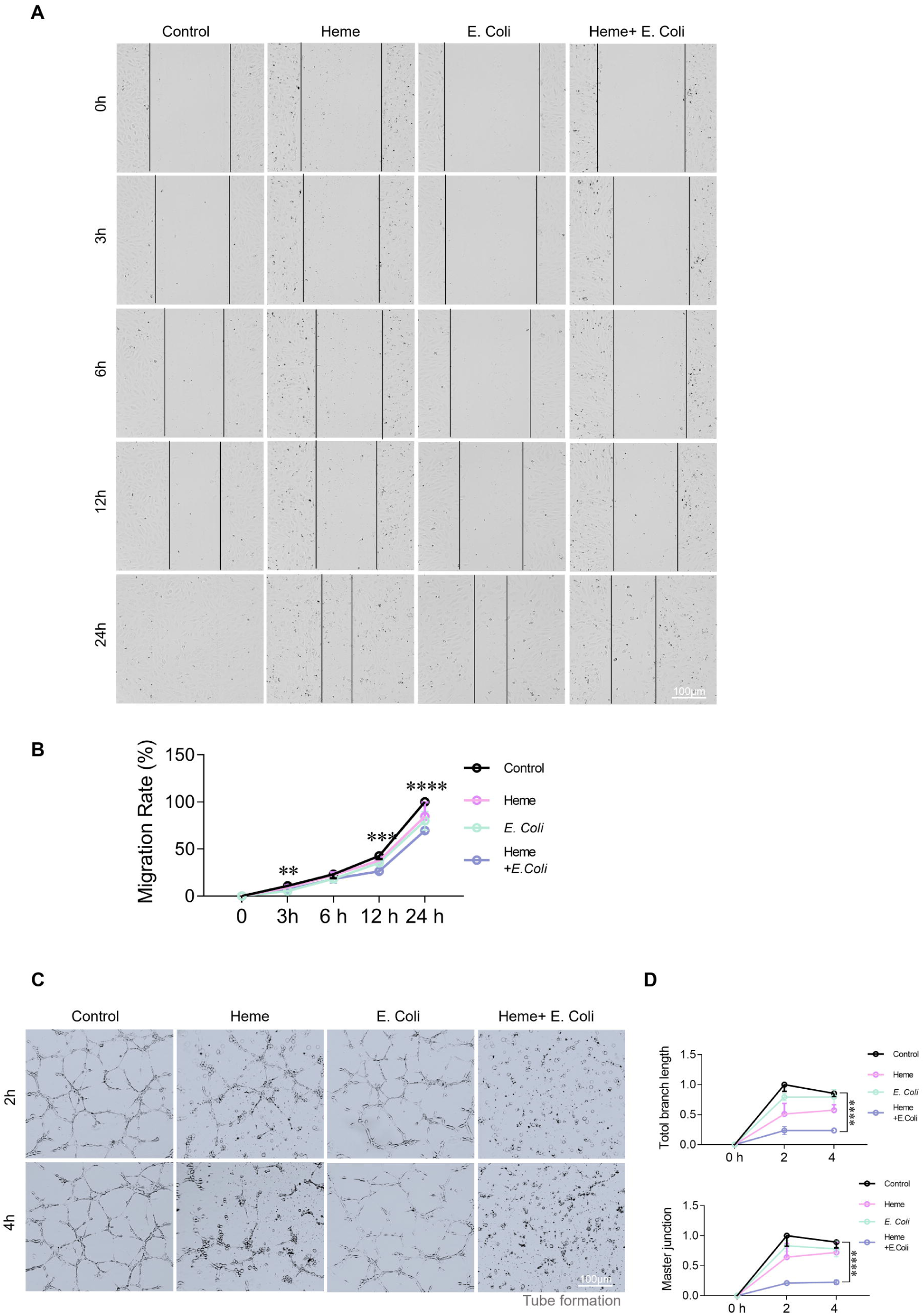
**Heme exacerbates bacteria-induced impairment of endothelial reparative capacity**. (**A, B**) Endothelial cell migration assay in HUVECs treated with heme, heat-killed *E. coli*, or both (n=6/group). (**C**) Endothelial tube formation assay under the same treatment conditions. (**D**) Quantification of total tube length and branching points in HUVECs (n=5/group). All data are presented as mean ± SD. Statistical analysis was performed using the unpaired two-tailed Student’s t-test: **PL<L0.01, ***PL<L0.001, ****PL<L0.0001.

### 4. Heme exacerbates bacterial induced endothelial senescence

To determine whether heme directly contributes to endothelial senescence under septic conditions, and to mimic the *in vivo* environment of bacterial infection in the presence of elevated heme, we treated HUVECs with heme (10µM), heat killed bacterial (*E.coli*, MOI:10), or a combination of both for 6 and 24 hours. Notably, treatment with either heme or *E.coli* alone induced only mild endothelial senescence, as indicated by β-gal staining (*Figures 4A-B*) and increased expression of senescence markers p16, p21, and acetyl-p53 (*Figures 4C-D*). Specifically, heme treatment led to 12.6% senescence, while E. coli exposure resulted in 10.6% senescence in HUVECs. Strikingly, co-treatment with heme and E. coli resulted in a dramatic increase in cellular senescence, reaching 76.5% in HUVECs as shown by β-gal staining (*Figures 4A-B*). Consistently, p21 immunostaining also revealed a robust increase in endothelial senescence in the co-treatment group compared to heme or E.coli alone (*Figures 4E-F*). Additionally, the combination of heme and E.coli significantly upregulated the expression of senescence-associated secretory phenotype (SASP) factors, including CXCL9, IL-α, MMP8, MMP9, ,CXCL2, FN1, and THBS1 compared to either treatment alone(*Figure 4I*). Effective vascular repair following injury is critical for cardiac functional recovery. To further assess whether endothelial senescence impairs this repair capacity, we evaluated endothelial cell migration, proliferation, and tube formation. Co-treatment with heme and E. coli significantly impaired all three functions—migration, proliferation, and tube formation, compared to single treatments or controls (*Figures 4G-H, S3A-C*). These findings suggest that heme not only directly promotes endothelial senescence but also acts synergistically with bacterial components to amplify cellular senescence in endothelial cells, thereby exacerbating vascular dysfunction and impairing vascular repair capacity during sepsis. This severe cardiac endothelial senescence may also help explain why many septic patients experience long-term cardiomyopathy.

**Figure. 4.**
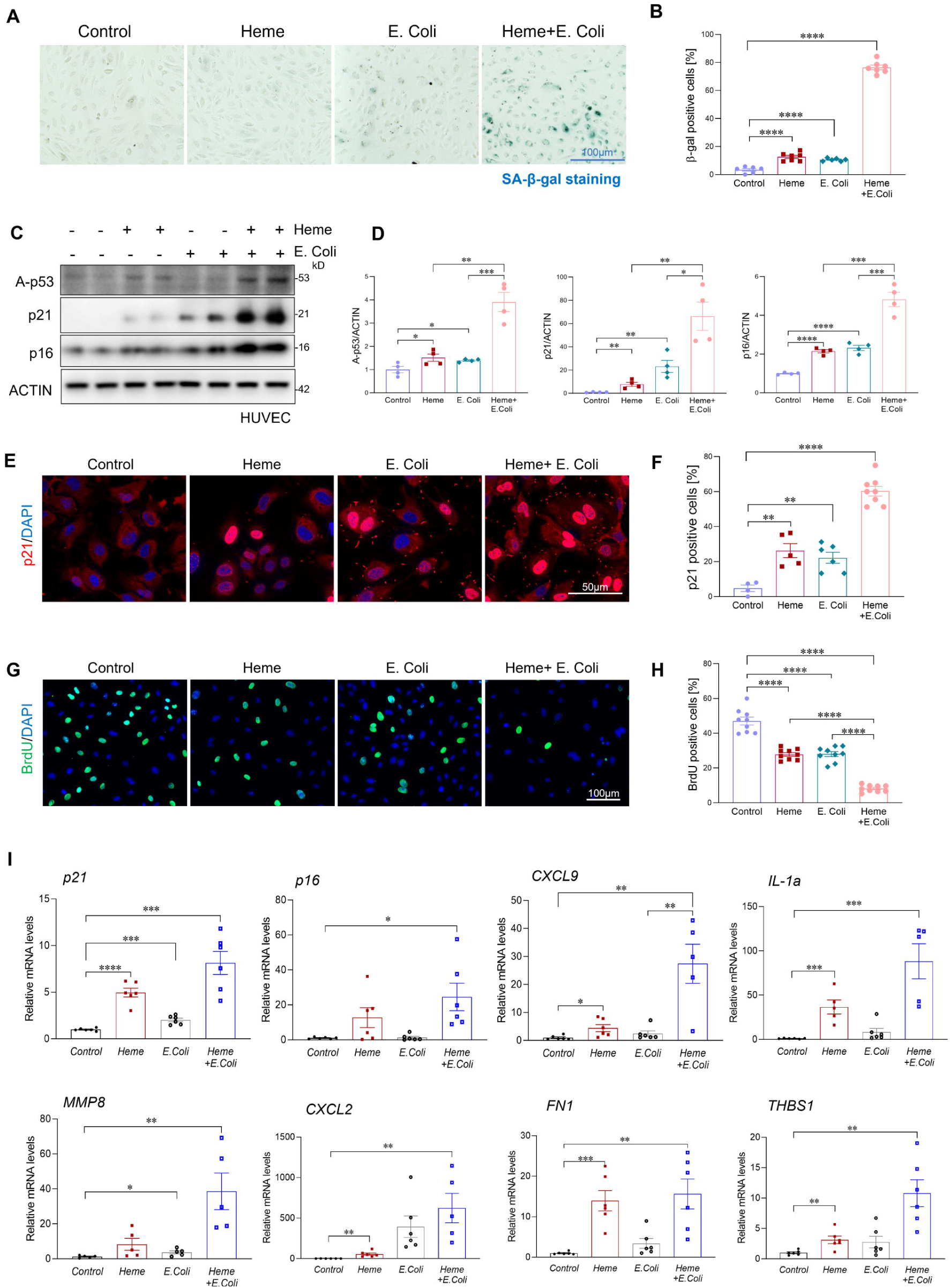
**Heme exacerbates bacterial induced endothelial senescence**. (**A, B**) β-galactosidase staining and quantification of HUVECs treated with heme (10LµM), heat-killed *E. coli* (MOI:10), or both for 24 hours (n=6/group). Scale bars, 100Lμm. (**C, D**) Western blot analysis and quantification of senescence markers p16, p21, and acetylated p53 in the indicated HUVEC groups (n=4/group). (**E, F**) Immunofluorescent staining and quantification of p21 in the indicated HUVEC groups (n=6/group). Scale bars, 50Lμm. (**G, H**) BrdU incorporation assay and quantification in the indicated HUVEC groups (n=6/group). Scale bars, 100Lμm. (**I**) qRT-PCR analysis of senescence-associated genes in the indicated HUVEC groups (n=6/group). All data are presented as mean ± SD. Statistical analysis was performed using the unpaired two-tailed Student’s t-test: *PL<L0.05, **PL<L0.01, ***PL<L0.001, ****PL<L0.0001.

### 5. Heme aggravates bacterial infection induced STING activation

To elucidate the mechanisms linking heme and bacterial induced endothelial senescence, we examined the activation of the cGAS-STING pathway, a known driver of cellular senescence^22, 30, 32^. As shown in Figure 5A-B, treatment of HUVECs with either heme or heat-killed E. coli alone resulted in only modest activation of the cGAS–STING pathway, as indicated by increased phosphorylation of STING, TBK1, and IRF3. However, co-treatment with heme and *E. coli* resulted in a marked enhancement of this pathway activation, with phosphorylation levels of STING, TBK1, and IRF3 increased by 2.7, 2.9, and 2.1 fold, respectively, compared to *E. coli* treatment alone (*Figures 5A-B*). Consistently, in vivo studies also revealed significantly enhanced activation of the cGAS–STING pathway in septic heart tissues following heme administration, compared to control septic heart tissues (*Figures 5H-I*).

**Figure. 5.**
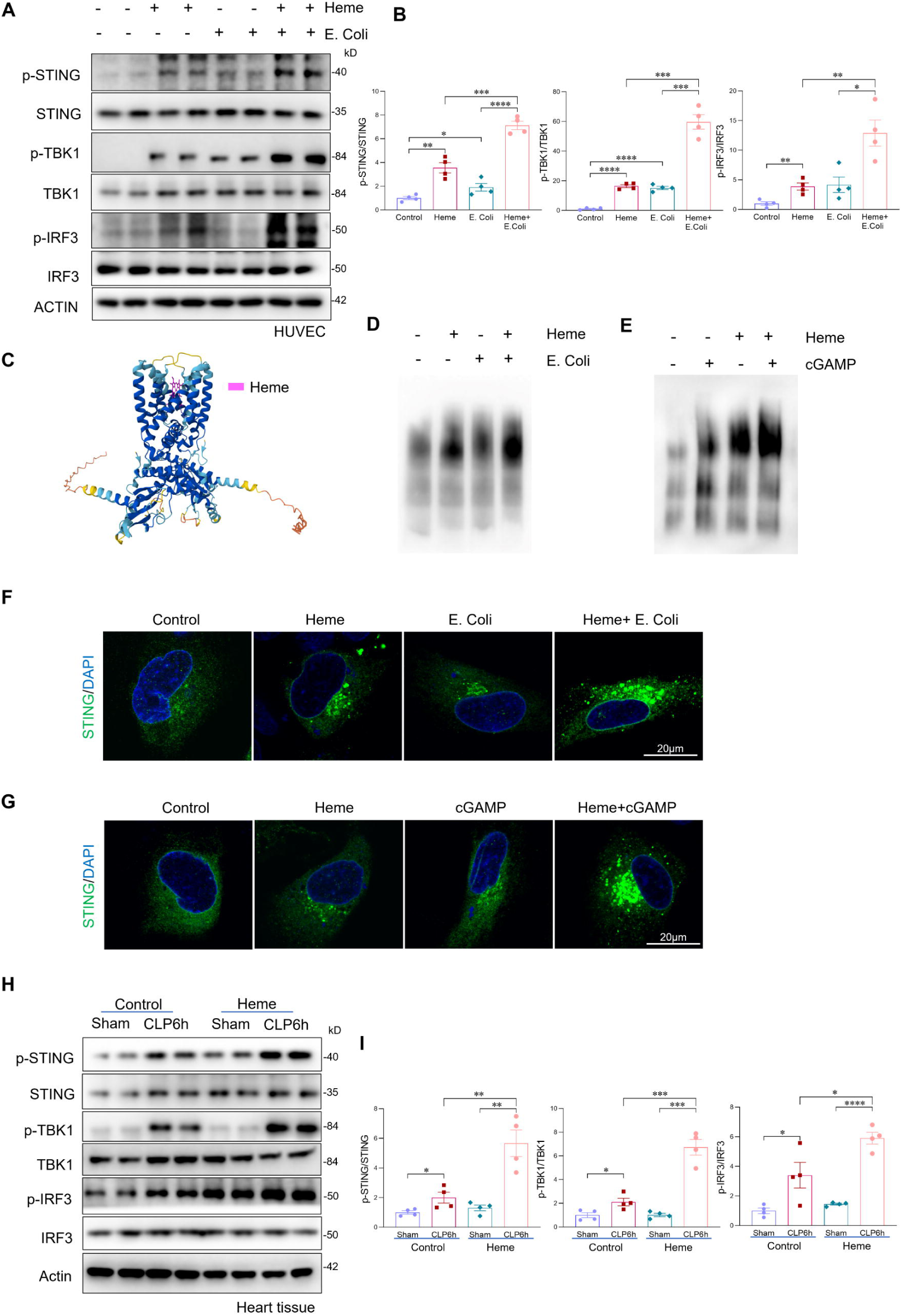
**Heme aggravates bacterial infection induced STING activation**. (**A, B**) Western blot analysis and quantification of p-STING/STING, p-TBK1/TBK1, and p-IRF3/IRF3 in HUVECs treated with heme (10LµM), heat-killed *E. coli* (MOI:10), or both for 6 hours (n=6/group). (**C**) Structural modeling of heme (pink) binding to the STING dimer complex using AlphaFold. (**D, E**) Native gel Western blot analysis of STING oligomerization in HUVECs treated with heme, *E. coli*, cGAMP, or their combinations. (**F, G**) Immunofluorescence staining of STING aggregates in HUVECs transfected with STING-HA and TBK1-Flag plasmids, followed by treatment with heme, *E. coli*, cGAMP, or their combinations. Scale bars, 20Lμm. (**H, I**) Western blot analysis and quantification of p-STING/STING, p-TBK1/TBK1, and p-IRF3/IRF3 in heart tissues from sham and septic mice treated with heme (n=4/group). All data are presented as mean ± SD. Statistical significance was determined using the unpaired two-tailed Student’s t-test: *PL<L0.05, **PL<L0.01, ***PL<L0.001, ****PL<L0.0001.

This robust activation of cGAS–STING signaling was not merely due to additive effects of heme and E. coli, but rather suggested a potential synergistic mechanism. To investigate whether heme directly binds to STING and facilitates its oligomerization, a critical step required for downstream activation of the pathway. We generate a three-dimensional model of the STING dimer in complex with heme. As show in Figure 5C, heme was predicated to bind to the STING dimer. Based on this, we hypothesized that heme binding promotes STING activation by enhancing its oligomerization during bacterial infection. To test this, we assessed STING oligomerization using native gel western blot analysis following treatment with heme, E. coli, or both. In untreated cells, STING predominantly existed in a dimeric form. Treatment with either heme or E. coli individually induced low levels of STING oligomer formation. However, combined treatment with heme and E. coli resulted in a substantial increase in high-order STING oligomers (*Figure 5D*), indicating enhanced oligomerization and synergistic pathway activation. To further validate this mechanism, we treated HUVECs with the canonical STING ligand cGAMP in the presence or absence of heme. Consistent with our hypothesis, co-treatment with cGAMP and heme induced significantly greater STING oligomerization than either agent alone (*Figure 5E*). In parallel, immunofluorescence staining revealed increased STING aggregation, particularly in cells co-treated with heme and E. coli or heme and cGAMP, supporting the formation of STING complexes (*Figures 5F-G & S4G-H*). To determine whether heme induced STING oligomerization enhances its interaction with TBK1, we transfected HUVECs with STING-HA and TBK-Flag plasmids and assessed their interaction by co-immunoprecipitation. As shown in Figures S4A–D, co-treatment with heme and *E. coli or* heme and *cGAMP* markedly enhanced the STING–TBK1 interaction compared to individual treatments. Collectively, these findings suggest that heme amplifies bacterial infection–induced activation of the cGAS–STING pathway by promoting STING oligomerization, thereby exacerbating endothelial cell senescence in sepsis.

**Figure S4.**
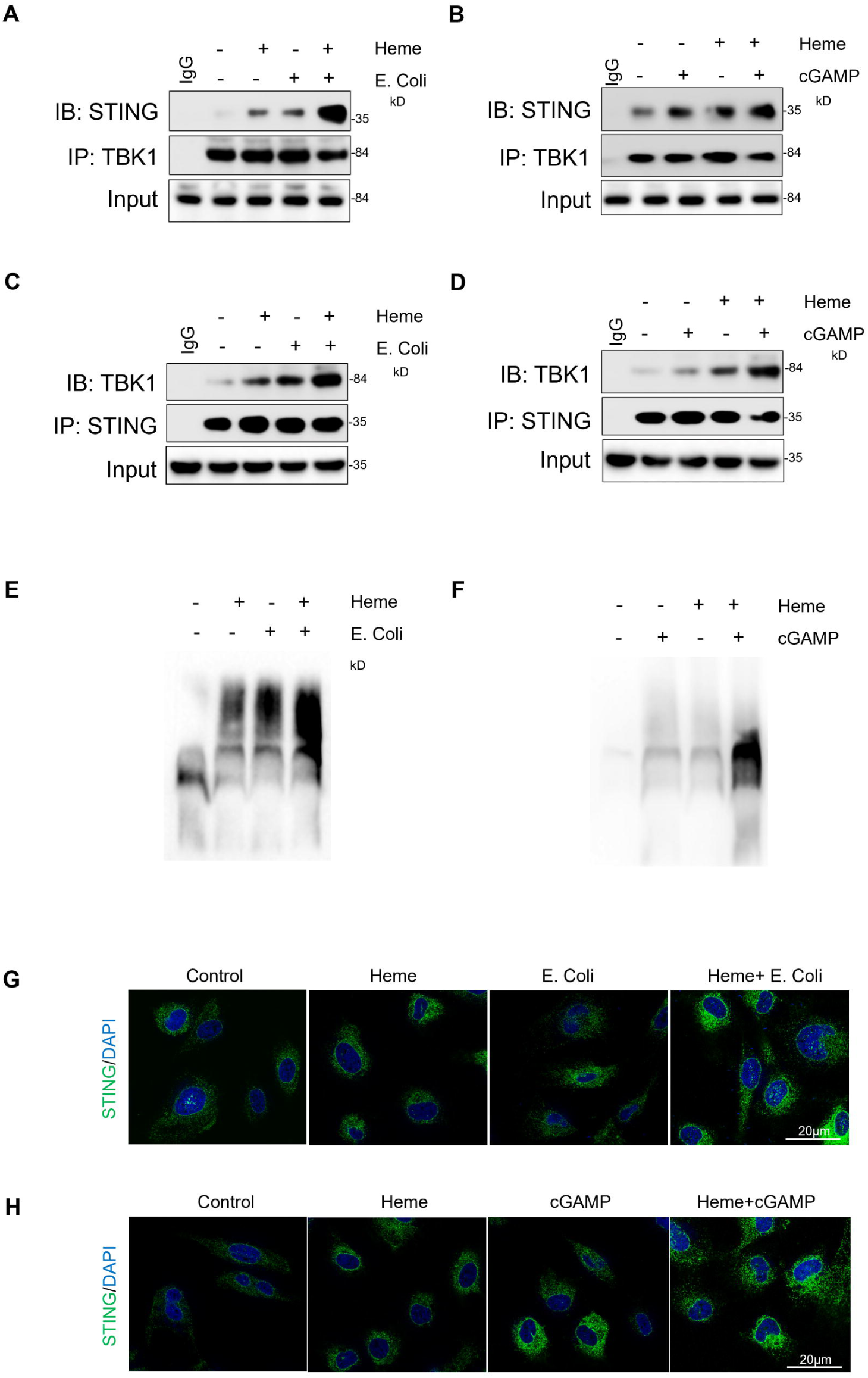
**Heme aggravates bacterial infection induced STING activation**. (**A-D**) Co-immunoprecipitation (Co-IP) assay indicate interaction between TBK1 and STING in HUVECs transfected with STING-HA and TBK1-Flag plasmids, followed by treatment with heme (10LµM), heat-killed *E. coli* (MOI:10), cGAMP (5Lµg/mL), or their combinations for 1 hour. (**E, F**) Native PAGE Western blot analysis demonstrating STING oligomerization in HUVECs transfected with STING-HA and TBK1-Flag plasmids under the same treatment conditions. (**G, H**) Immunofluorescence staining of STING aggregates in HUVECs following treatment with heme, *E. coli*, cGAMP, or combinations. Scale bars, 20Lμm. All data are presented as mean ± SD. Statistical significance was determined using the unpaired two-tailed Student’s t-test: *PL<L0.05, **PL<L0.01, ***PL<L0.001, ****PL<L0.0001.

### 6. STING activation contributes to sepsis induced cardiac endothelial senescence

We next investigated the role of cGAS–STING signaling in sepsis-induced cardiac endothelial senescence. Mice were treated with a STING inhibitor (C-176) or vehicle control immediately following CLP-induced sepsis. Heart tissues were collected 24 hours post-CLP. STING inhibition significantly reduced endothelial senescence in cardiac tissue, as evidenced by decreased p21 and p16 expression compared to vehicle-treated controls (*Figures 6A-B, S5A-B).* Moreover, echocardiographic analysis revealed that pharmacological inhibition of STING markedly improved cardiac function in septic mice, as indicated by increased ejection fraction (EF) and fractional shortening (FS) (*Figures 6C&6E-F*). To further determine whether long-term STING inhibition promotes sustained cardiac recovery, we treated septic mice with C-176 or vehicle every other day for two weeks. Long-term STING inhibition significantly reduced cardiac endothelial senescence and improved cardiac functional recovery (*Figure 6E-F&S5C-F*).

**Figure. 6.**
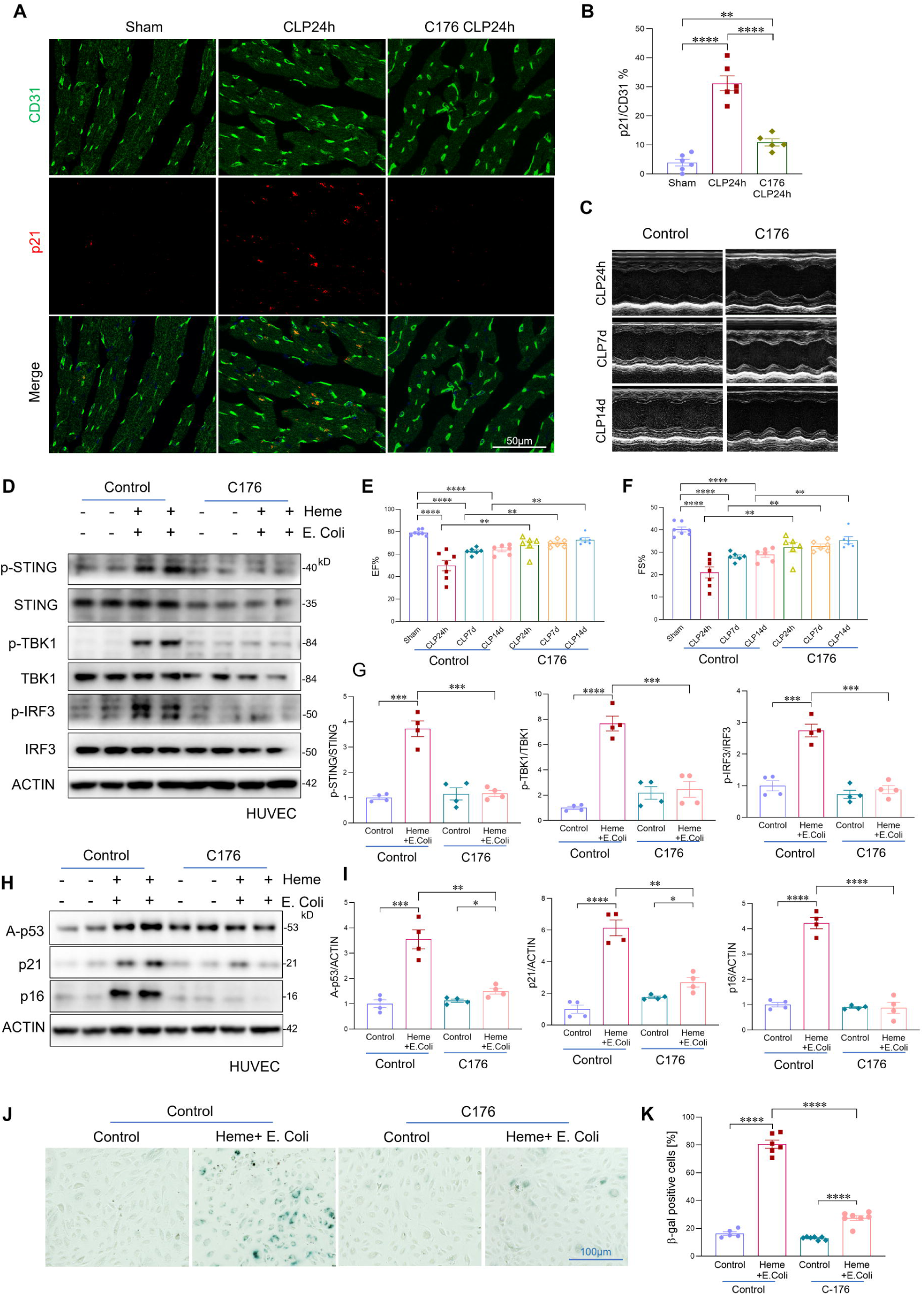
**STING activation contributes to sepsis induced cardiac endothelial senescence**. (**A, B**) Co-immunofluorescent staining and quantification of the senescence marker p21 with the endothelial cell marker CD31 in cardiac tissues from sham and septic mice treated with a STING inhibitor or vehicle control at 24 hours post-CLP (n=6 mice/group). Scale bars, 50Lμm. (**C**) Representative echocardiographic images assessing ejection fraction (EF) and fractional shortening (FS) at 24 hours, 7 days, and 14 days post-CLP in mice treated with vehicle or STING inhibitor. (**D**) Western blot analysis of p-STING/STING, p-TBK1/TBK1, and p-IRF3/IRF3 in HUVECs treated with heme + heat-killed *E. coli*, with or without the STING inhibitor C-176 (2.5LμM) (n=6/group). (**E, F**) Echocardiographic assessment of EF and FS in septic mice treated with STING inhibitor or vehicle control at 24 hours, 7 days, and 14 days post-CLP, as shown in (C) (n=6 mice/group). (**G**) Quantification of Western blot analysis of p-STING/STING, p-TBK1/TBK1, and p-IRF3/IRF3 in the indicated HUVEC groups, as shown in (**D**) (n=4/group). (**H, I**) Western blot analysis and quantification of senescence markers p16, p21, and acetylated p53 in the indicated HUVEC groups (n=6/group). (**J, K**) β-galactosidase (β-gal) staining and quantification of HUVECs treated with combined heme and *E. coli* with or without the STING inhibitor C-176 (n=6/group). Scale bars, 100Lμm. All data are presented as mean ± SD. Statistical analysis was performed using the unpaired two-tailed Student’s t-test: *PL<L0.05, **PL<L0.01, ***PL<L0.001, ****PL<L0.0001.

**Figure S5.**
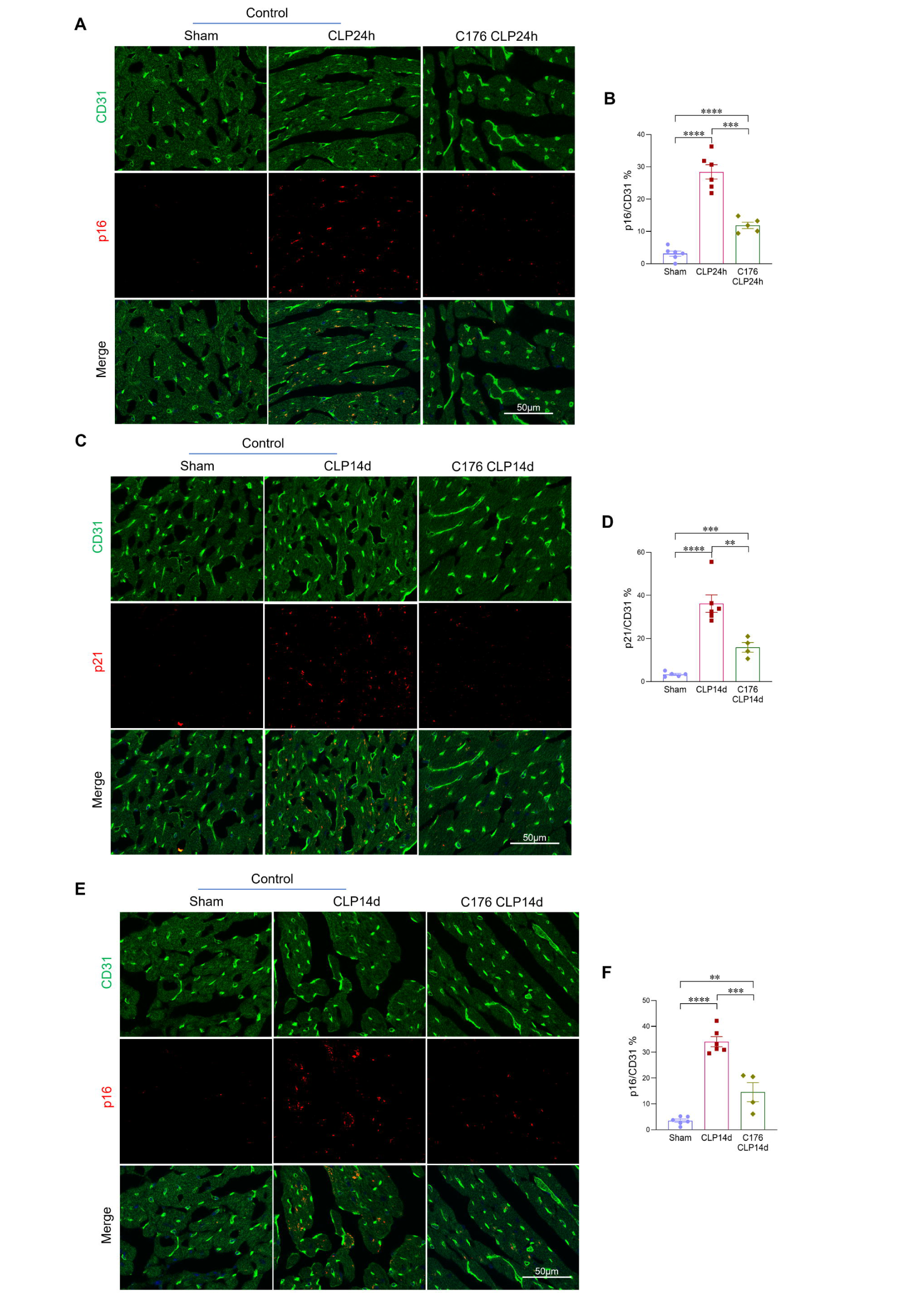
**STING inhibition attenuates sepsis-induced cardiac endothelial senescence**. (**A, B**) Immunofluorescent co-staining and quantification of the senescence marker p16 with the endothelial marker CD31 in heart tissues from sham and septic mice treated with a STING inhibitor or vehicle at 24 hours post-CLP (n=5-6 mice/group). Scale bars, 50Lμm. (**C–E**) Immunofluorescent co-staining and quantification of senescence markers p16 or p21 with the endothelial marker CD31 in heart tissues from sham and septic mice treated with a STING inhibitor or vehicle at 14 days post-CLP (n=4-6 mice/group). Scale bars, 50Lμm. All data are presented as mean ± SD. Statistical analysis was performed using the unpaired two-tailed Student’s t-test: **PL<L0.01, ***PL<L0.001, ****PL<L0.0001.

To assess whether STING activation directly contributes to heme and bacterial induced endothelial senescence, we treated HUVECs with heme combined with E. coli, or with septic plasma, in the presence or absence of a STING inhibitor. STING inhibition significantly suppressed activation of the cGAS–STING pathway in response to heme–bacterial challenge, as shown by reduced phosphorylation of STING, TBK1, and IRF3 (*Figures 6D&6G).* In parallel, STING inhibition markedly attenuated endothelial senescence induced by either heme plus *E.coli*, or septic plasma, as indicated by reduced β-galactosidase activity (*Figures 6J-K&S6A-B*). Consistent with these findings, STING inhibition reduced immunostaining of p21 (*Figures S6C-D*) and lowered protein expression of senescence markers p21, p16, and acetylated p53 (*Figures 6H-I*). Additionally, STING inhibition mitigated the detrimental effects of heme and E. coli on endothelial reparative function, significantly restoring endothelial proliferation, and tube formation capacity (*Figures S6E-H*). Collectively, these findings demonstrate that cGAS–STING signaling plays a critical role in driving endothelial senescence and cardiac dysfunction during sepsis. Inhibition of STING enhances endothelial repair capacity and promotes cardiac functional recovery, underscoring its therapeutic potential in sepsis-associated cardiovascular complications.

**Figure S6.**
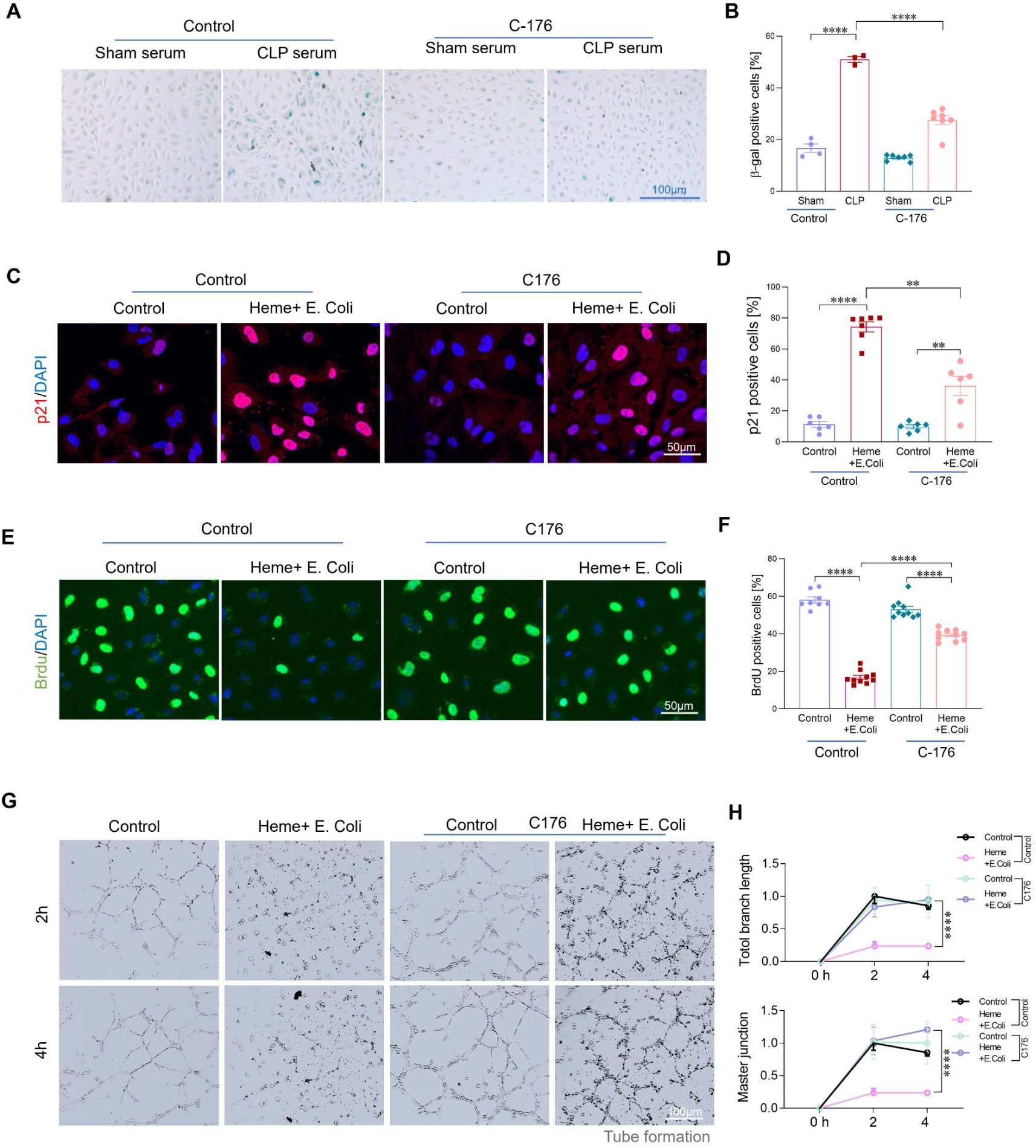
**STING inhibition restores endothelial reparative function in vitro**. (**A, B**) β-galactosidase staining and quantification of HUVECs treated with septic plasma, with or without STING inhibitor (C-176, 2.5LµM). (**C, D**) Immunofluorescent staining and quantification of senescence marker p21 in HUVECs treated with septic plasma, with or without STING inhibitor (n=6/group). Scale bars, 50Lμm. (**E–H**) Endothelial proliferation (BrdU) and tube formation assays following treatment with combined heme and E. coli, with or without STING inhibitor. All data are presented as mean ± SD. Statistical analysis was performed using the unpaired two-tailed Student’s t-test: *PL<L0.05, **PL<L0.01, ****PL<L0.0001.

### 7. Hemopexin upregulation alleviates sepsis-induced endothelial senescence and cardiac dysfunction

Hemopexin (HPX), a liver derived heme scavenger, plays a critical in neutralizing excess free heme and preventing its cytotoxic effects^22, 23^. Given our earlier findings that circulating free heme levels were markedly elevated in non-survivors compared to survivors, we next examined HPX expression across these groups. Notably, HPX levels were significantly higher in survivors than non-survivors (*Figures 7A-C*), suggesting a protective role of HPX during sepsis. To determine whether enhancing HPX expression could attenuate sepsis induced endothelial senescence and cardiac dysfunction, we administered AAV-HPX (1×10¹¹GC/mice) to overexpress HPX, with AAV-Con as a control. Two weeks after AAV delivery, the mice were subjected to CLP sepsis. ELISA assay confirmed successful elevated HPX levels in plasma (*Figure 7D*). Following sepsis induction, mice treated with AAV–HPX exhibited significantly reduced circulating free heme levels (*Figure 7E*). Increased HPX expression attenuated cardiac endothelial senescence, as evidenced by decreased expression of p21 and p16 at both 24 hours and 14 days post-sepsis (*Figures 7F-G, S7A-D&S8C-D*). Consistently, Ki67 staining showed that endothelial cell proliferative capacity was enhanced in septic mice with increased HPX expression, compared to control septic mice (*Figures S8A-B*). Furthermore, AAV–HPX treatment improved cardiac functional recovery, as indicated by increased ejection fraction (EF) and fractional shortening (FS) on echocardiographic analysis at 24 hours, 7 days, and 14 days following sepsis (*Figures 7H-J*). Collectively, these findings demonstrate that enhancing heme clearance through increased HPX expression mitigates sepsis-induced cardiac endothelial senescence and promotes cardiac functional recovery.

**Figure. 7.**
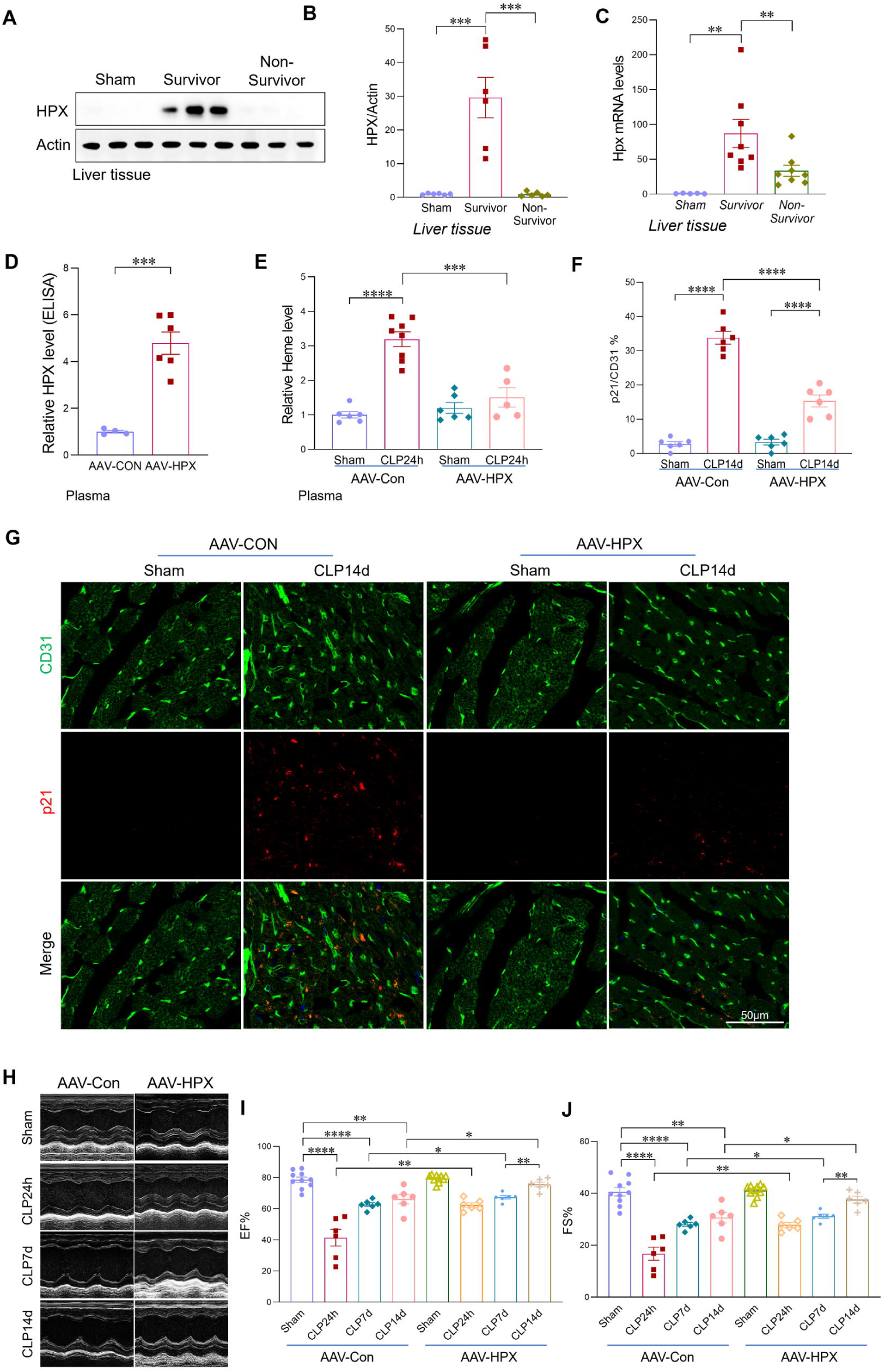
**Hemopexin upregulation alleviates sepsis-induced endothelial senescence and cardiac dysfunction**. (**A, B**) Western blot analysis and quantification of HPX expression in liver tissues from sham mice, septic survivors, and septic non-survivors at 24 hours post-CLP (n=6/group). (**C)** qRT-PCR analysis of HPX expression in liver tissues from sham, septic survivors, and septic non-survivors (n=6-8/group). (**D**) ELISA confirmation of HPX overexpression in plasma following AAV–HPX administration (1×10¹¹ GC/mouse), with AAV–Con as the control (n=6/group). (**E**) Plasma free heme levels measured 24 hours post-CLP in AAV–HPX and AAV–control mice (n=6-8/group). (**F, G**) Co-immunofluorescent staining and quantification of the senescence marker p21 with the endothelial cell marker CD31 in heart tissues from the indicated groups. Scale bars, 50Lμm. (**H–J**) Echocardiographic assessment of ejection fraction (EF) and fractional shortening (FS) in the indicated groups at 24 hours, 7 days, and 14 days post-sepsis (n=6-8/group). All data are presented as mean ± SD. Statistical analysis was performed using the unpaired two-tailed Student’s t-test: *PL<L0.05, **PL<L0.01, ***PL<L0.001, ****PL<L0.0001.

**Figure S7.**
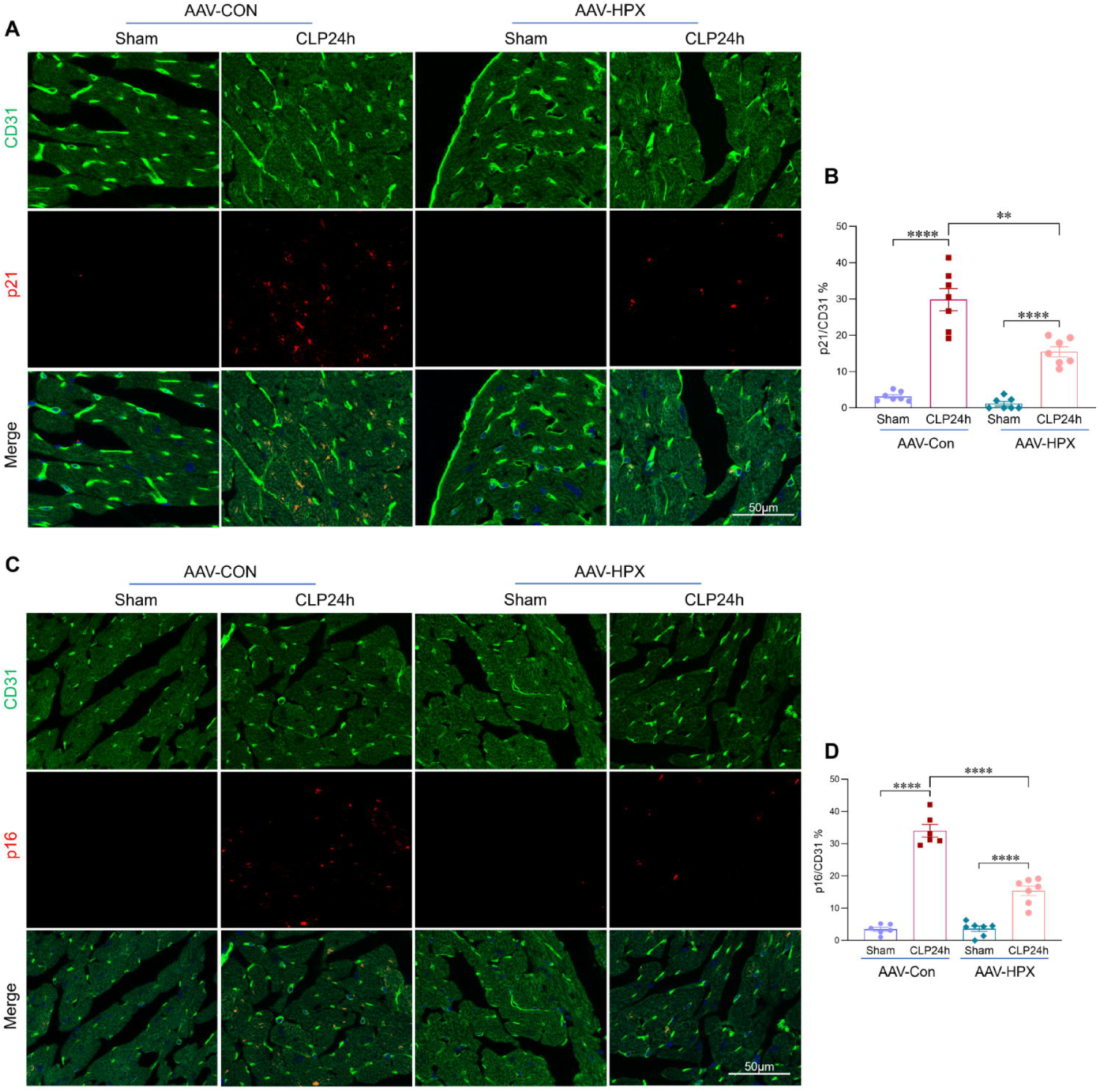
**Increased hemopexin expression attenuates sepsis-induced cardiac endothelial senescence**. (**A–D**) Immunostaining and quantification of the senescence markers p21 or p16 with endothelial cell markers in sham and septic heart tissues from AAV-HPX and AAV-Con mice at 24 hours post-CLP (n=6-7/group). Scale bars, 50Lμm. All data are presented as mean ± SD. Statistical analysis was performed using the unpaired two-tailed Student’s t-test: **PL<L0.01, ****PL<L0.0001.

**Figure S8.**
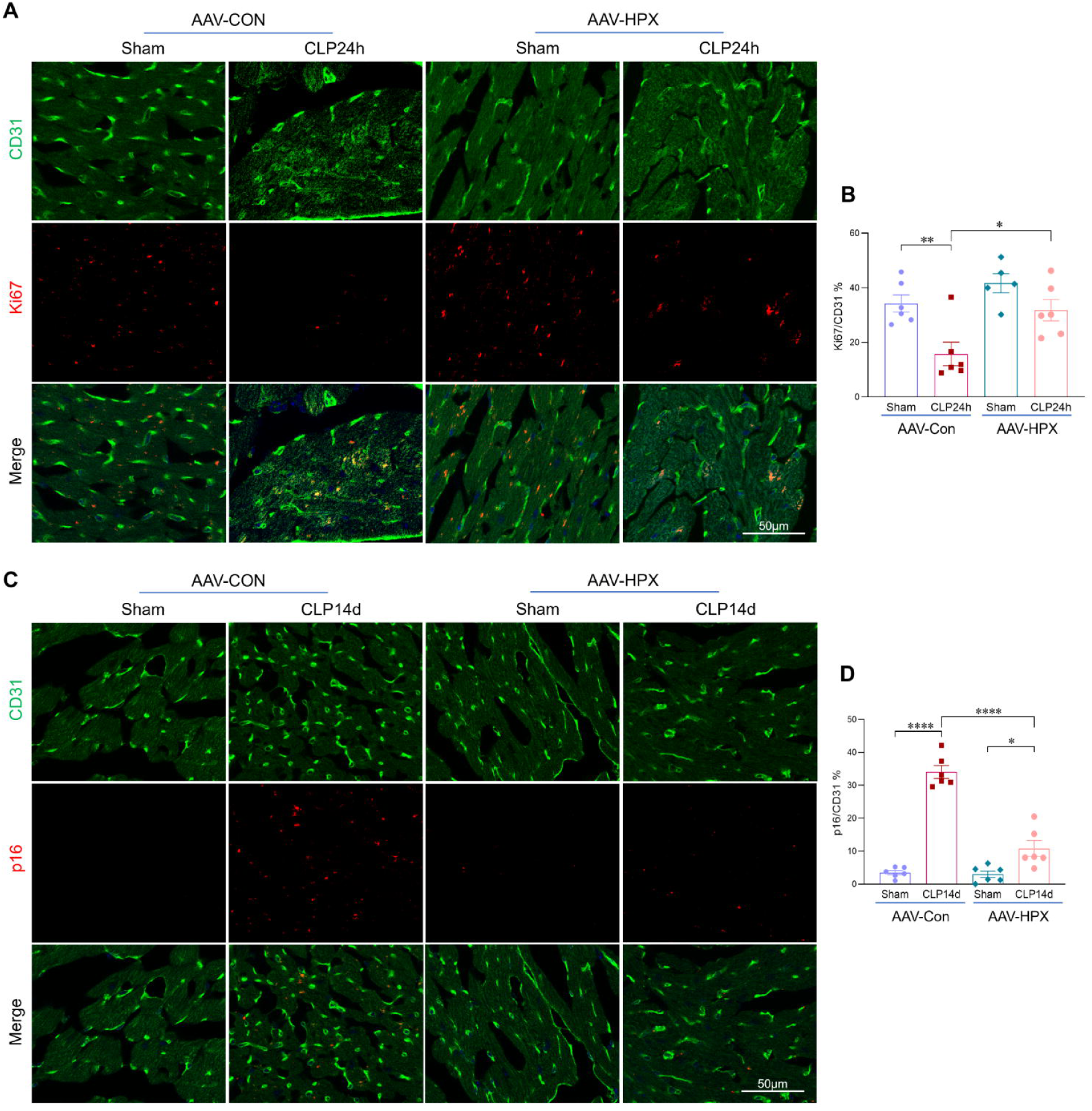
**Increased hemopexin expression attenuates sepsis-induced cardiac endothelial senescence and enhances endothelial proliferative capacity**. (**A, B**) Co-immunostaining and quantification of the cellular proliferation marker Ki67 with the endothelial marker CD31 in sham and septic heart tissues from AAV–HPX and AAV–Con mice at 24 hours post-sepsis. Scale bars, 50Lμm. (**C, D**) Co-immunostaining and quantification of the senescence marker p16 with the endothelial marker CD31 in heart tissues from AAV–HPX and AAV–Con mice at 14 days post-sepsis. Scale bars, 50Lμm. All data are presented as mean ± SD. Statistical analysis was performed using the unpaired two-tailed Student’s t-test: *PL<L0.05, **PL<L0.01, ****PL<L0.0001.

## Discussion

Sepsis-induced cardiac dysfunction is a life-threatening complication that substantially contributes to sepsis-related mortality^3, 4, 5^. Despite its clinical significance, the underlying cellular and molecular mechanisms remain incompletely understood. In this study, we identify cardiac endothelial senescence as a key pathological feature of sepsis and demonstrate that elevated levels of free heme, a byproduct of hemolysis, plays a pivotal role in driving this response. Our findings reveal a novel role for heme in activating the cGAS-STING pathway and promoting endothelial senescence, thereby impairing vascular repair and contributing to cardiac dysfunction during sepsis (*Figure 8*). These results highlight both STING inhibition and heme clearance as promising therapeutic strategies for preserving cardiovascular function in septic patients.

**Figure. 8.**
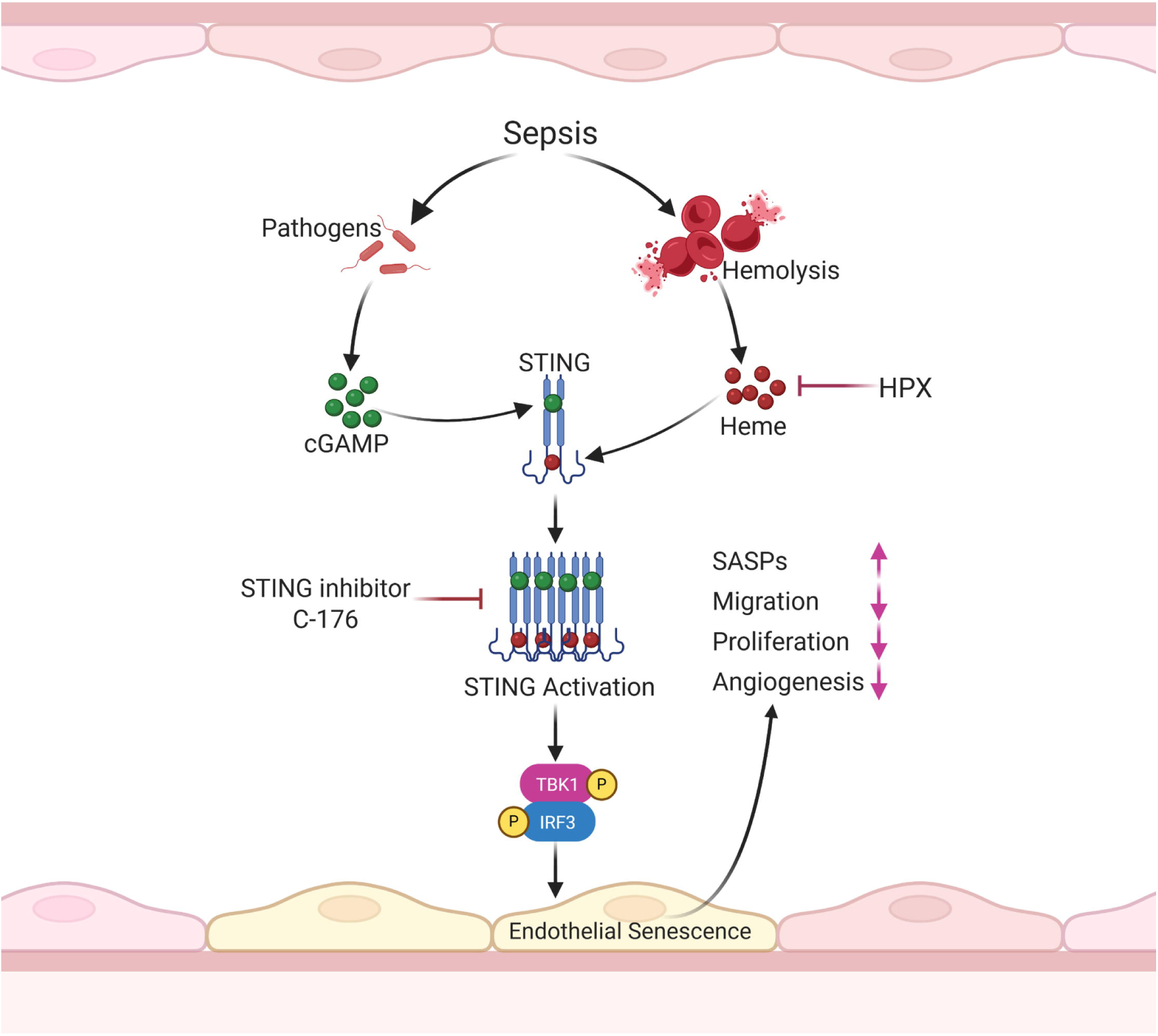
**Schematic Illustration**. Heme drives cardiac endothelial senescence in sepsis through STING activation. During sepsis, free heme—a byproduct of hemolysis—acts as a novel ligand for STING, amplifying bacterial infection–induced STING polymerization and downstream signaling activation, thereby promoting endothelial senescence. Senescent endothelial cells contribute to sepsis-induced cardiac dysfunction by upregulating senescence-associated secretory phenotype (SASP) factors and impairing vascular repair through diminished endothelial proliferation, migration, and angiogenesis. Pharmacological inhibition of STING or hemopexin (HPX)-mediated heme clearance attenuates endothelial senescence and improves cardiac functional recovery under septic conditions.

Endothelial senescence, a hallmark of aging, contributes to the pathogenesis of several cardiovascular diseases (CVDs), including stroke, atherosclerosis, and hypertension^15, 16, 17, 19, 20, 21, 36^. However, whether endothelial senescence occurs in the context of sepsis, and whether senescent endothelial cells contribute to sepsis-associated cardiac dysfunction, has not been previously explored. In this study, we show for the first time that cellular senescence is markedly elevated in the heart during sepsis, with endothelial cells representing the predominant senescent population. Notably, the extent of cardiac endothelial senescence correlated with disease severity and was significantly greater in non-surviving animals. Senescent endothelial cells exhibited impaired proliferative, angiogenic, and migratory capacity, which compromise vascular repair and hinder cardiac functional recovery following sepsis. These findings may explain why patients recovering from sepsis remain at increased risk for long-term cardiovascular complications.

A major advance of this study is the identification of free heme as a key driver of cardiac endothelial senescence. We demonstrate that circulating free heme levels correlate with the severity of endothelial senescence and cardiac dysfunction in septic mice. Moreover, administration of exogenous heme exacerbated both senescence and cardiac impairment, confirming its causative role in this process. One of the strengths of this study is the discovery that heme amplifies bacterial infection induced activation of the cGAS–STING pathway, a known driver of cellular senescence^22, 30, 32,37^. Co-treatment with heme and bacterial components synergistically enhanced cGAS–STING signaling beyond additive effects. A previous study showed that compound C53^38^, a small-molecule agonist, binds to a cryptic pocket within the transmembrane domain of STING, located between the two subunits of the STING dimer. In the presence of cGAMP, this interaction induces robust STING oligomerization and activation^38^. Interestingly, using protein–ligand binding prediction through AlphaFold, we found that heme is also bind within the cryptic pocket of the STING transmembrane domain. This observation led us to hypothesize that heme might act similarly to C53 in promoting STING activation by facilitating its oligomerization. As anticipated, the presence of heme significantly enhanced STING oligomerization and activation in response to bacterial challenge. This effect was further validated by co-treatment with cGAMP, confirming that heme augments STING signaling through oligomerization. These results provide compelling evidence that the interaction between pathogen-and damage-associated signals orchestrates a maladaptive endothelial senescence response during sepsis.

In our in vivo model, pharmacological inhibition of STING significantly alleviated endothelial senescence and improved cardiac functional recovery. Previous studies have established that activation of the cGAS–STING pathway during sepsis leads to excessive production of proinflammatory cytokines, contributing to widespread tissue injury^39, 40, 41^. Given that sustained inflammation is a known inducer of cellular senescence^42, 43^, we investigated whether cGAS– STING signaling directly contributes to endothelial senescence under septic conditions. To address this, we conducted in vitro experiments using cardiac endothelial cells and found that STING inhibition markedly reduced heme-and bacteria-induced endothelial senescence. These findings suggest that, beyond its established role in promoting inflammation, cGAS–STING signaling is a key driver of cardiac endothelial senescence and dysfunction during sepsis.

Hemopexin (HPX), a liver-derived heme-binding protein, functions as a scavenger of free heme^22, 23, 25^. Previous clinical studies have shown that septic patients with lower hemopexin levels tend to experience earlier mortality compared to those with higher levels. In our study, we similarly found that HPX levels were significantly higher in survivors than in non-survivors.

Notably, increasing HPX expression significantly reduced circulating free heme levels, attenuated cardiac endothelial senescence, and restored cardiac function in septic mice. These findings support the concept that enhancing heme clearance represents a viable strategy to mitigate organ dysfunction in sepsis.

Overall, our study has several important implications. First, it highlights cardiac endothelial senescence as a pathological hallmark and therapeutic target in septic cardiomyopathy. Second, it identifies free heme as a pathogenic factor that integrates hemolysis and immune activation to exacerbate endothelial damage and cardiac injury. Finally, it supports the therapeutic potential of targeting the cGAS–STING pathway or enhancing heme clearance to reduce vascular dysfunction and improve cardiac functional recovery.

There are limitations to consider. First, although we focused on cardiac endothelial senescence, senescence in other organs such as the lungs and kidneys were not examined and may also contribute to systemic dysfunction during sepsis. Second, the predicted interaction between heme and STING was based on computational modeling using AlphaFold. Further structural validation is required to confirm this binding and its functional consequences.

In conclusion, our study identifies a novel heme–STING–senescence axis that drives cardiac endothelial dysfunction during sepsis. By linking hemolysis to vascular injury and cardiac impairment, these findings provide a compelling rationale for therapeutically targeting heme clearance and STING signaling to preserve cardiovascular integrity and improve outcomes in sepsis.

## Materials and Methods

### Animals and Housing

Wild-type C57BL/6 mice were obtained from Jackson Laboratory and housed at the Institutional Animal Care and Use Facility at East Tennessee State University. Male and female mice aged 3-5 months were categorized as young, while those aged 22-24 months were classified as aged. All animal care and experimental procedures were approved by the ETSU Committee on Animal Care and Use and adhered to NIH guidelines to ensure humane treatment.

### Sepsis Model: Cecal Ligation and Puncture (CLP)

Polymicrobial sepsis was induced using the cecal ligation and puncture (CLP) model as described previously. Briefly, mice were anesthetized with isoflurane and positioned on a heated surgical pad. Following a midline incision, approximately one-third of the cecum was ligated and punctured once using a 23-gauge needle. After repositioning the cecum, the abdominal cavity was closed. The peritoneal wall was sutured with sterile 4-0 Dafilon sutures and the skin was closed with a surgical staple (Autoclip 9 mm; Cat:12020-09). Sham-operated mice underwent similar procedures without ligation or puncture. Following CLP, mice were administered a single subcutaneous dose of resuscitative fluid. Liver tissues and plasma were collected at 6 and 24 hours post CLP for immunostaining, heme level quantification, and bacterial colony analysis.

### Cell culture and treatment

Human umbilical vein endothelial cells (HUVECs) were obtained from the American Type Culture Collection (ATCC, PCS-100-013) and cultured in Vascular Cell Basal Medium (ATCC, PCS-100-030) supplemented with the Endothelial Cell Growth Kit–VEGF (ATCC, PCS-100-041). Cells were maintained at 37°C in a humidified incubator with 5% CO₂. HUVECs at passages below P6 were considered young, while those at passage P12 or higher were classified as aged. For treatment experiments, HUVECs were exposed to the following stimuli and inhibitors, either alone or in combination as indicated: heme (10 μM, Hemin, Sigma, Cat: 51280-5G), heat-killed E. coli (MOI: 20, ATCC, Cat: BAA197), STING inhibitor (2.5µM, C-176, MCE, Cat: HY-112906). Cells were treated for the indicated times to assess proliferation using Brdu staining, mitigation using wound healing assay, and angiogenic capacity using the tube formation assay. Cell senescence was determined via β-galactosidase (SA-β-Gal) activity using the Senescence β-Galactosidase Staining Kit (CST, Cat: 9860S) following the manufacturers’ protocols. Additionally, the expression of senescence markers was analyzed by qPCR assay, Western blot and immunostaining.

### Adeno-Associated Virus (AAV) packaging and in vivo Administration

Recombinant adeno-associated virus (AAV) vectors expressing the human hemopexin (HPX) gene under a CMV promoter (pAAV-CMV-HPX, ABM, Cat: 23884101) were generated using a triple-plasmid system, including a packaging plasmid (pAAV2/8, Addgene Cat: 112864) and a helper plasmid (pAdDeltaF6, Addgene Cat: 112867). The pAAV-CMV-Blank vector served as a negative control. HEK293T cells were co-transfected with these plasmids using Lipofectamine 3000, and viral particles were harvested 72 hours later. HEK293T cells were co-transfected with the plasmids using Lipofectamine 3000, and viral particles were harvested 72 hours post-transfection. Viral purification and titering were performed according to published protocols. Mice received intravenous injections of AAV-HPX or control AAV at a single dose of 1 × 10¹¹ genome copies (GC) in 100 µL of sterile PBS via the tail vein. Two weeks post-AAV delivery, the cecal ligation and puncture (CLP) model of sepsis was performed as previously described. Transfection efficiency was confirmed by assessing HPX expression in liver tissues.

### Immunofluorescence, confocal imaging, and analysis

At the indicated time points following CLP-induced sepsis, both sham and septic mice were anesthetized, and liver tissues were collected and fixed in 4% paraformaldehyde (PFA) for 12 hours at 4 °C, followed by immersion in 30% sucrose overnight. Samples were then embedded in OCT compound and sectioned at 10 μm thickness using a Leica CM3050S cryostat. For in vitro studies, HUVECs were cultured on chamber slides, washed with PBS, and fixed in 2% PFA for 20 minutes at room temperature. Both cryosections and HUVECs were washed three times with PBS, permeabilized with 0.5% Triton X-100 in PBS for 20 minutes at room temperature, and then blocked with 10% goat serum for 1 hour. Samples were incubated overnight at 4°C with the following primary antibodies: anti-CD31 (endothelial marker, R&D, Cat #AF3628, 1:100), anti-p16 Cell Signaling Technology, Cat# #80772S, 1:100), anti-p21 (Cell Signaling Technology, Cat #2947S, 1:100), and anti-STING (ABclonal, Cat# A21051, 1:100). Following primary incubation, sections and cells were washed once with PBST and twice with PBS, then incubated with appropriate Alexa Fluor–conjugated secondary antibodies for 1 hour at room temperature. After three final PBST washes, samples were mounted with DAPI-containing fluorescence mounting medium (Thermo Fisher Scientific, Cat# P36981) to stain nuclei.

Confocal images were captured using a Leica TCS SP8 microscope and processed using ImageJ software (NIH) for quantitative and qualitative analysis.

### Protein Extraction and Western Blot Analysis

Protein extraction from heart tissues and in vitro cultured HUVECs was performed using RIPA lysis buffer (Thermo Fisher Scientific, Cat: 89900) supplemented with protease and phosphatase inhibitors (Sigma-Aldrich, Cat: PPC2020). Protein concentrations were measured using a BCA protein assay kit (Thermo Fisher Scientific, Cat: 23223). For blue native PAGE (BN-PAGE), protein samples were prepared using the NativePAGE^™^ Sample Prep Kit (Invitrogen, Cat# BN2005) containing 10%DDM, along with NativePAGE^™^ 4× Sample Buffer (Invitrogen, Cat# BN2003). The 5% Coomassie G-250 sample additive (Invitrogen, Cat# BN2002) was added immediately prior to loading. Samples were resolved on NativePAGE^™^ Novex® 3–12% Bis-Tris gels (Invitrogen, Cat# NP0321BOX), and electrophoresis was performed using the XCell™ SureLock^™^ Mini-Cell system with the recommended Dark Blue and Light Blue cathode buffers. Following migration of the dye front to one-third of the gel, Dark Blue buffer was replaced with Light Blue buffer to reduce background interference during Western transfer. After electrophoresis, proteins were transferred to PVDF membranes using the XCell II^™^ Blot Module and NuPAGE® Transfer Buffer (Thermo Fisher Scientific, Cat# 88518, NP00061). Equal amounts of protein were separated by electrophoresis on Bis-Tris protein gels and transferred onto nitrocellulose membranes. Membranes were blocked with 5% BSA in TBS containing 0.5% Tween-20 (TBST) for 1 hour at room temperature, followed by incubation with specific primary antibodies overnight at 4°C. After washing with TBST, membranes were incubated with HRP-conjugated secondary antibodies (Cell Signaling Technology, Cat: 7074) for 1 hour at room temperature, followed by additional washes with TBST to remove unbound antibodies. Bands were visualized using the SuperSignal™ West Femto Maximum Sensitivity Substrate (Thermo Fisher Scientific, Cat: 34096), optimized for detecting low-abundance targets, and SuperSignal™ West Pico PLUS Chemiluminescent Substrate (Thermo Fisher Scientific, Cat: 34580) for higher-abundance targets. Imaging was conducted with the G:Box Chemi gel documentation system (GeneSys Version: 1.8.10.0), and band intensities were quantified using Image J software.

### Plasmid Transfection of STING and TBK1

To increase the expression of STING and TBK1 in HUVECs, cells were seeded at ∼60% confluence and transfected with either the human STING1 ORF cDNA expression plasmid containing a C-terminal HA tag or the human TBK1 ORF mammalian expression plasmid with a C-terminal Flag tag (Sino Biological, Cat# HG29810-CY and HG11023-CF, respectively). Transfection was performed using polyethylenimine (PEI, linear, MW 25,000, Polysciences, Cat# 23966) at a DNA:PEI ratio of 1:3. Briefly, plasmid DNA and PEI were each diluted in Opti-MEM (Gibco, Cat# 31985070), incubated at room temperature for 15 minutes to allow complex formation, and then added dropwise to the cells. After 4 hours, the transfection medium was replaced with complete endothelial growth medium. Cells were harvested 48 hours post-transfection for downstream applications.

### Antibodies

The following primary and secondary antibodies were used in this study: Anti-STING (Cat: 13647S), anti-p-STING (Cat: 72971S), anti-TBK1 (Cat: 3504S), anti-p-TBK1 (Cat: 5483S), anti-IRF3 (Cat: 4302S), anti-p-IRF3 (Cat: 4947S), anti-p-IRF3 (Cat: 4947S), anti-p-IRF3 (Cat: 4947S), p21 (Cat: 2947S) were purchased from Cell Signaling Technology. Anti-p16 (Cat: A8571), Anti-AP53 (Cat: A19836), Anti-Hemopexin (HPX) (Cat: A5603), Anti-DDDDK-Tag (Cat: AE005) were purchased from Abclonal.

### RNA-seq and data analysis

Total RNA was extracted from sham and septic mouse heart tissues using RNAzol RT reagent (Molecular Research Center, Cat# RN 190) according to the manufacturer’s protocol. RNA quality and concentration were assessed using an Agilent Bioanalyzer 2100, and samples with RNA integrity number (RIN) >L6.0 were used for library preparation. RNA quality control, library preparation, sequencing, and data analysis were performed by Genewiz (Azenta Life Sciences). Differential expression analysis was conducted using standard bioinformatics pipelines provided by the service provider. Gene Set Enrichment Analysis (GSEA) was performed using the GSEA software (Broad Institute) to assess pathway-level changes.

### Quantitative RT-PCR Analysis

Total RNA was isolated from mouse heart tissues and cultured cells using RNAzol RT reagent (Molecular Research Center, Cat: RN 190) according to the manufacturer’s protocol. cDNA was synthesized from 0.1–1 μg of total RNA using the High-Capacity cDNA Reverse Transcription Kit (Thermo Fisher Scientific, Cat: 4368814). Quantitative real-time PCR (qRT-PCR) was performed using SYBR Select Master Mix on a Bio-Rad CFX96 real-time PCR detection system. The following PCR primers were used: H-CDKN1A (forward, 5’-TGTCCGTCAGAACCCATGC-3’; reverse, 5’-AAAGTCGAAGTTCCATCGCTC-3’), M-Cdkn1a (forward, 5’-CCTGGTGATGTCCGACCTG-3’; reverse, 5’-CCATGAGCGCATCGCAATC-3’), H-CDKN2A (forward, 5’-GGAGGCCGATCCAGGTCAT-3’; reverse, 5’-ATGGTTACTGCCTCTGGTGC-3’), M-Cdkn2a (forward, 5’-CGCAGGTTCTTGGTCACTGT-3’; reverse, 5’-TGTTCACGAAAGCCAGAGCG-3’), M-Cxcl2 (forward, 5’-ATTCTGTGACCATCCCCTCAT-3’; reverse, 5’-TGTATGTGCCTCTGAACCCAC-3’), H-CXCL2 (forward, 5’-AGATCAATGTGACGGCAGGG-3’; reverse, 5’-TCTCTGCTCTAACACAGAGGGA-3’), H-CXCL1 (forward, 5’-TGCTGCCACTAATGCTGATGT-3’; reverse, 5’-CTCAGGAACCAATCTTTGCACT-3’), M-Il1a (forward, 5’-CGAAGACTACAGTTCTGCCATT-3’; reverse, 5’-GACGTTTCAGAGGTTCTCAGAG-3’), H-IL1A (forward, 5’-TGGTAGTAGCAACCAACGGGA-3’; reverse, 5’-ACTTTGATTGAGGGCGTCATTC-3’), H-SERPINE1 (forward, 5’-CCTGGGCACTTACAGGAAGG-3’; reverse, 5’-GGTCCGATTCGTCGTCAAATAAC-3’), H-FN1 (forward, 5’-CGGTGGCTGTCAGTCAAAG-3’; reverse, 5’-AAACCTCGGCTTCCTCCATAA-3’), H-Thbs1 (forward, 5’-AGACTCCGCATCGCAAAGG-3’; reverse, 5’-TCACCACGTTGTTGTCAAGGG-3’), M-Mmp8 (forward, 5’-TCTTCCTCCACACACAGCTTG-3’; reverse, 5’-CTGCAACCATCGTGGCATTC-3’), H-MMP8 (forward, 5’-TGCTCTTACTCCATGTGCAGA-3’; reverse, 5’-TCCAGGTAGTCCTGAACAGTTT-3’).

### Measurement of Heme Levels in plasma

Blood samples were collected via cardiac puncture into tubes containing heparin and centrifuged at 3000g for 20 minutes to separate plasma. Plasma heme levels were measured using the Heme Assay Kit (Sigma-Aldrich, Cat:MAK316) according to the manufacturer’s instructions. Plasma hemopexin (HPX) levels were quantified using an ELISA kit (Abclonal, Cat: RK09237)

### Echocardiography

M-mode tracings were used to measure left ventricular (LV) wall thickness, LV end-systolic diameter (LVESD), and LV end-diastolic diameter (LVEDD). Percent fractional shortening (%FS) and ejection fraction (EF) were calculated as previously described^44, 45^.

### Statistical Analysis

All experiments were performed at least three independent times and representative data are shown. Statistical analysis between two groups were performed by an unpaired two-tailed Student’s t-test, while one-way ANOVA with Tukey’s post hoc test was used for multiple comparisons. A p-value less than 0.05 was considered statistically significant. Data are expressed as mean ± SD. Analyses were performed using GraphPad Prism 8.4.3 software.

## Acknowledgements

We thank the ETSU Molecular Biology Core Facility (MBCF) for providing G:BOX imaging, Nanodrop, and Primer ordering services, and the ETSU Microscopy Core Facility for confocal imaging support. We also thank Dr. Krishna Singh for providing access to the Vevo 1100 ultrasound system from VisualSonics (Fujifilm).

## Author Contributions

TL designed and performed experiments, analyzed data, interpreted results, and wrote the original manuscript. PZ, TZ, JA, FT, GC performed experiments and contributed to data analysis and interpretation. XZ, LL, DW, and CL assisted with experimental design and data interpretation, reviewed, and revised the manuscript. XW designed and supervised the project and wrote the manuscript. All authors reviewed and approved the final manuscript.

## Data availability

All data are available in the main text or the supplementary materials. All original data generated or analyzed during this study are available from the corresponding author upon request.

## Ethics approval

All animal studies were approved by the Animal Care and Use Committee at East Tennessee State University and were conducted in accordance with institutional guidelines and relevant regulations.

